# Molecular sorting of nitrogenase catalytic cofactors

**DOI:** 10.1101/2025.01.21.634024

**Authors:** Alvaro Salinero-Lanzarote, Josh Lian, Gil Namkoong, Daniel L. M. Suess, Luis M. Rubio, Dennis R. Dean, Ana Pérez-González

**Author notes:** Corresponding authors**: Ana Pérez-González, Dennis R. Dean,**. These authors contributed equally to this manuscript (Alvaro Salinero-Lanzarote, Josh Lian, Gil Namkoong).

## Abstract

The free-living diazotroph *Azotobacter vinelandii* produces three genetically distinct but functionally and mechanistically similar nitrogenase isozymes, designated as Mo-dependent, V-dependent, and Fe-only. They respectively harbor nearly identical catalytic cofactors that are distinguished by a heterometal site occupied by Mo (FeMo-cofactor), V (FeV-cofactor), or Fe (FeFe-cofactor). Completion of FeMo-cofactor and FeV-cofactor formation occurs on molecular scaffolds prior to delivery to their catalytic partners. In contrast, completion of FeFe-cofactor assembly occurs directly within its cognate catalytic partner. Because hybrid nitrogenase species that contain the incorrect cofactor type cannot reduce N_2_ to support diazotrophic growth there must be a way to prevent misincorporation of an incorrect cofactor when different nitrogenase isozyme systems are produced at the same time. Here, we show that fidelity of the Fe-only nitrogenase is preserved by blocking the misincorporation of either FeMo-cofactor or FeV-cofactor during its maturation. This protection is accomplished by a two-domain protein, designated AnfO. It is shown that the N-terminal domain of AnfO binds to an immature form of the Fe-only nitrogenase and the C-terminal domain, tethered to the N-terminal domain by a flexible linker, has the capacity to capture FeMo- and FeV-cofactor. AnfO does not prevent the normal activation of Fe-only nitrogenase because completion of FeFe-cofactor assembly occurs within its catalytic partner and, therefore, is never available for capture by AnfO. These results support a post-translational mechanism involving the molecular sorting of structurally similar metallocofactors that involve both protein-protein interactions and metallocofactor binding while exploiting differential pathways for nitrogenase associated catalytic cofactor assembly.

## Introduction

The nitrogenases are the only known catalysts for biological nitrogen fixation, the enzymatic conversion of inert N_2_ to metabolically tractable NH_3_. The free-living diazotroph *Azotobacter vinelandii* has the capacity to produce three genetically distinct but structurally, functionally and mechanistically similar nitrogenase isoforms (1–3). All three isoforms are two-component systems that include a system-specific Fe protein (component II) involved in the ATP-dependent electron delivery to its catalytic partner (component I) that provides the site for substrate binding and reduction. The nitrogenase isoforms are designated Mo-dependent, V-dependent, and Fe-only based on the presence or absence of a heterometal (Mo, V, or Fe) contained within their corresponding catalytic cofactors. These cofactors are structurally similar, each having an Fe-S-C containing core with an apical heterometal atom coordinated to the organic constituent homocitrate. According to their metal compositions, the corresponding cofactors are denoted FeMo-cofactor, FeV-cofactor, and FeFe-cofactor, and their cognate component I proteins are designated MoFe protein, VFe protein and FeFe protein, respectively (4). In addition to the catalytic cofactors contained in the MoFe protein (products of *nifDK*), VFe protein (products of *vnfDGK*), and FeFe protein (products of *anfDGK*), each also contains an 8Fe-7S “P-cluster” believed to mediate electron transfer from a redox-active 4Fe-4S cluster contained within the isoform-specific Fe protein to the corresponding catalytic cofactor and, ultimately, substrate. Crystallographically determined structural models of the nitrogenase MoFe protein (5), VFe protein (6), and FeFe protein (7), as well as their associated metalloclusters, are shown in Fig. 1.

**Figure 1.**
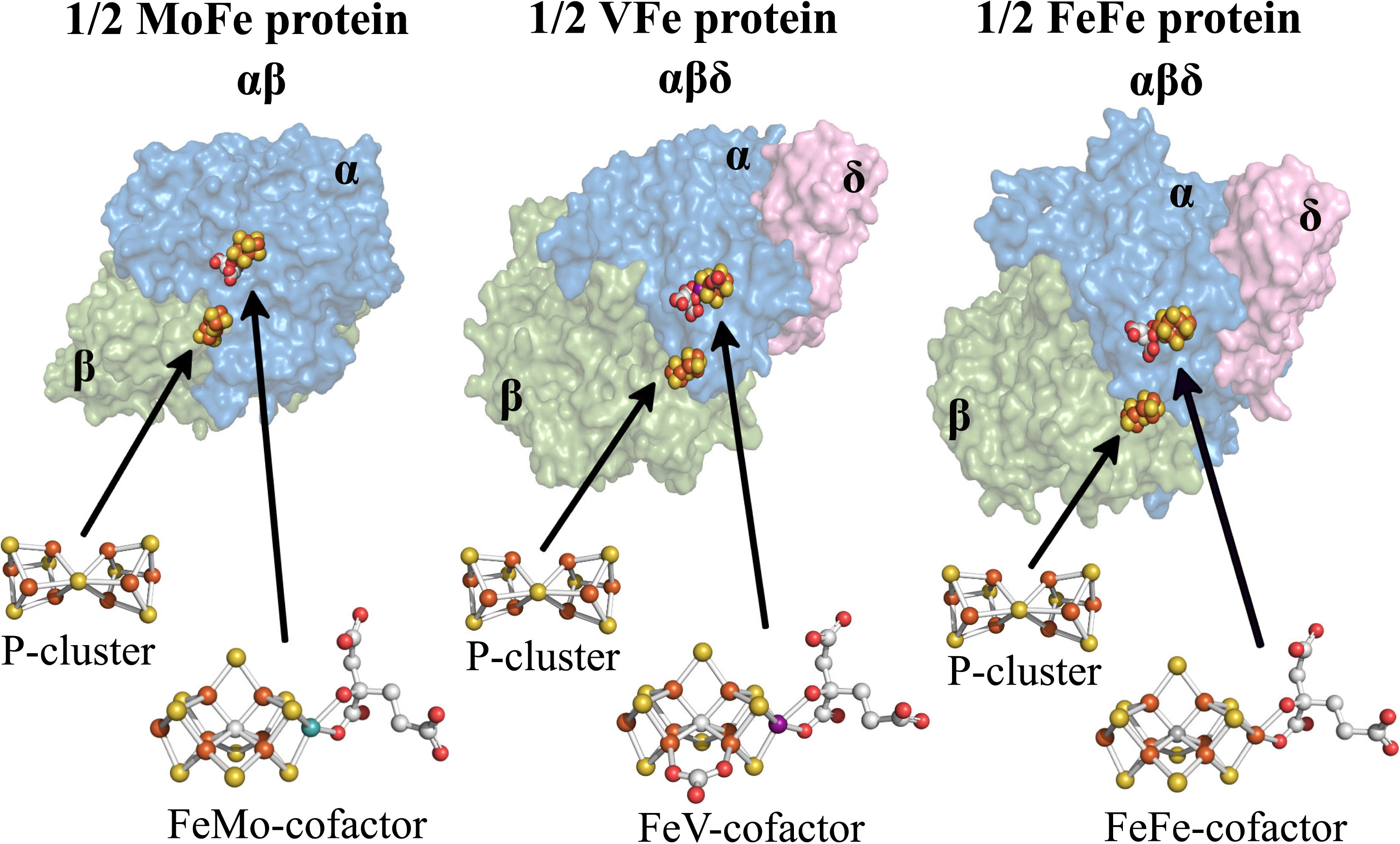
MoFe-VFe- and FeFe-protein structural models. MoFe protein is an ɑ_2_β_2_ heterotetramer whereas the VFe protein and FeFe protein are ɑ_2_β_2_δ_2_ heterohexamers. Only single αβ and αβ8 units of the respective proteins are shown with ɑ-subunits shaded blue, β-subunits shaded green, and δ-subunits shaded pink. α, β, and δ subunits are encoded by *anfD*, *anfK*, and *anfG*, respectively. Each ɑβ or ɑβδ unit contains a catalytic cofactor, respectively designated FeMo-cofactor, FeV-cofactor, or FeFe-cofactor, and an 8Fe-7S P cluster. Atoms in the cofactor structures are represented as; orange: iron, yellow: sulfur; grey: carbon; red: oxygen; turquoise: molybdenum; purple: vanadium. MoFe protein, FeMo-cofactor and P-cluster structures were extracted from PDB 3U7Q (5); VFe protein and FeV-cofactor from PDB 5N6Y (6); FeFe protein and FeFe-cofactor from PDB 8BOQ (7). Figures were generated using PyMol (58).

For all three isoforms, P-cluster assembly precedes insertion of the corresponding catalytic cofactors (8–10). An 8Fe-9S-C species, designated NifB-co or L-cluster by some investigators, is a common precursor for all three catalytic cofactors (2, 11). In the case of FeMo-cofactor formation, NifB-co is further processed on an α_2_β_2_ molecular scaffold designated NifEN, upon which an apical Fe atom within NifB-co is substituted by Mo and homocitrate attachment occurs (12–14). NifB-co is believed to be processed in the same way on a structurally equivalent α_2_β_2_ scaffold designated VnfEN (15, 16), which is involved in V insertion and homocitrate attachment to complete FeV-cofactor formation. In contrast, completion of FeFe-cofactor formation does not require an assembly scaffold (16). Instead, homocitrate attachment to NifB-co occurs directly within an immature form of the FeFe protein. These cofactor assembly processes are schematically shown in Fig. 2.

**Figure 2.**
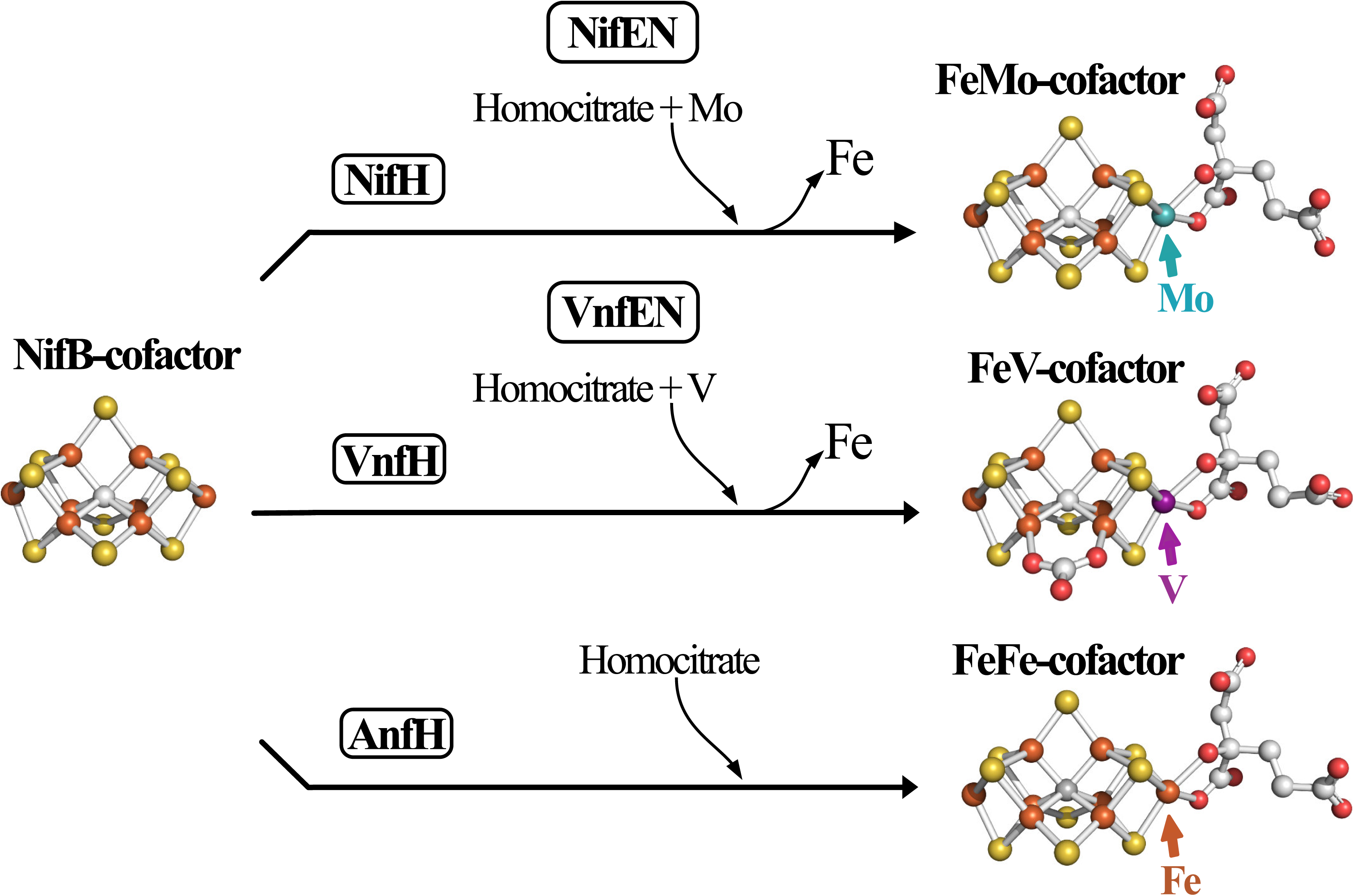
Schematic representation of active-site cofactors assembly. NifB-co is a common precursor for all three catalytic cofactors. The corresponding nitrogenase reductase (component II) for each system—NifH, VnfH, and AnfH—are believed to be required for NifB-co processing for each system (59), although this has not been experimentally established in the case of FeFe-cofactor formation (60). Completion of FeMo-cofactor and FeV-cofactor formation respectively occur on the NifEN and VnfEN scaffolds upon which an apical Fe atom within NifB-co is replaced by Mo or V and the organic constituent homocitrate is attached (11–13). The role of VnfEN in FeV-cofactor formation has not been established experimentally but is inferred by a requirement for VnfEN to form FeV-cofactor and primary structure similarities between NifEN and VnfEN (15). No molecular scaffold is required for FeFe-cofactor formation (16). Atoms in the cofactor structures are represented as orange: iron; yellow: sulfur; grey: carbon; red: oxygen; turquoise: molybdenum; purple: vanadium. FeMo-cofactor was extracted from PDB 3U7Q (5); FeV-cofactor from PDB 5N6Y (6); FeFe-cofactor from PDB 8BOQ (7). NifB-co structure is derived from FeFe-cofactor lacking homocitrate. Figures were generated using PyMol (58).

Despite their similar structures and cofactors, nitrogenase isozymes are not equal in enzymatic efficiency; the Mo-dependent nitrogenase is the most efficient catalyst for N_2_ reduction, whereas the Fe-only nitrogenase is the least efficient (3). This disparity in catalytic efficiency provides the impetus for exquisite control of the differential accumulation of nitrogenase isoforms in response to the physiological demand for fixed nitrogen as well as the bioavailability of the heterometals. Such regulation, which mainly occurs at the transcriptional level, involves the global nitrogen regulatory elements, system-specific regulatory elements, and Mo availability-modulated regulatory elements (17). Nevertheless, the physiological transition from conditions that favor accumulation of one nitrogenase isoform to a different isoform could result in the incorporation of the incorrect catalytic cofactor, rendering the hybrid species inactive and, therefore, delaying adjustment to the new environment.

Based on the physiological and biochemical phenotypes resulting from inactivation of the *A. vinelandii anfO* gene associated with the Fe-only nitrogenase, the existence of a post-translational mechanism that controls the fidelity of FeFe protein maturation by preventing incorporation of the incorrect cofactor was proposed (18). Namely, it was found that when *anfO* is inactivated and V is available in the growth medium, even at trace levels, FeV-cofactor becomes inserted into the FeFe protein resulting in the accumulation of a hybrid species that cannot support N_2_ reduction. How AnfO accomplishes that function has not been determined, nor has its ability to prevent misincorporation of the FeMo-cofactor. In the present work, we provide evidence for a novel post-translational mechanism involving AnfO in the molecular sorting of structurally similar metallocofactors through both protein-protein interactions and metallocofactor binding.

## Results

### Strains

To experimentally evaluate the adventitious incorporation FeMo-cofactor into FeFe protein, a series of strains were isolated for which Mo repression of the Fe-only nitrogenase expression is attenuated. For the parental strain, designated DJ2560, and its derivatives, the structural genes *nifDK* encoding the MoFe protein and the *vnfDGK* genes encoding the VFe protein were inactivated to prevent their capacity to reduce N_2_, which could result in NH_3_-directed repression of *anfHDGK* encoding Fe-only nitrogenase accumulation (19). The *modE1* gene was also disabled to attenuate Mo repression of Fe-only nitrogenase gene expression. Deletion of *modE1* provided W-tolerant diazotrophic growth capacity, as already reported (16). For all strains, the *vnfEN* genes, encoding the scaffold for FeV-cofactor formation, were disabled to prevent formation of FeV-cofactor and, therefore, any possible FeV-cofactor associated misincorporation into the FeFe protein (18). Detailed descriptions of the genotypes of strains used are described in Table S1 and S2.

Evaluation of the possible role of AnfO in preventing the misincorporation of FeMo-cofactor into FeFe protein was also made possible by isolation of strains derived from DJ2560 that are either deleted for *anfO* (DJ2821) or deleted for both *anfO* and *nifEN* (DJ2831). As noted above, NifEN provides a scaffold for the completion of FeMo-cofactor biosynthesis and, therefore, strain DJ2831 cannot produce FeMo-cofactor.

### AnfO prevents misincorporation of FeMo-cofactor into the FeFe protein

Figure 3 shows the growth capacities of relevant strains used in the present work when cultured under various conditions. All strains grow when a fixed nitrogen source, ammonium acetate, is added to the culture medium. A control strain (DJ2240) disabled for all the three nitrogenase isoforms is unable to sustain diazotrophic growth under any condition. A strain deleted for *anfO* (DJ2821) but otherwise having an intact capacity for producing Fe-only nitrogenase exhibits no diazotrophic growth when cultured in the presence of 5 μM Mo. In contrast, a strain having an intact *anfO* (DJ2560) exhibits normal diazotrophic growth when cultured in the presence of 5 μM Mo. The Mo-sensitive growth phenotype of DJ2821 deleted for *anfO* is reversed when combined with a *nifEN* deletion to produce strain DJ2831. The growth phenotypes are consistent with an incapacitation of FeFe protein to support N_2_ reduction as a result of misincorporation of FeMo-cofactor to form a hybrid FeFe species when *anfO* is deleted.

**Figure 3.**
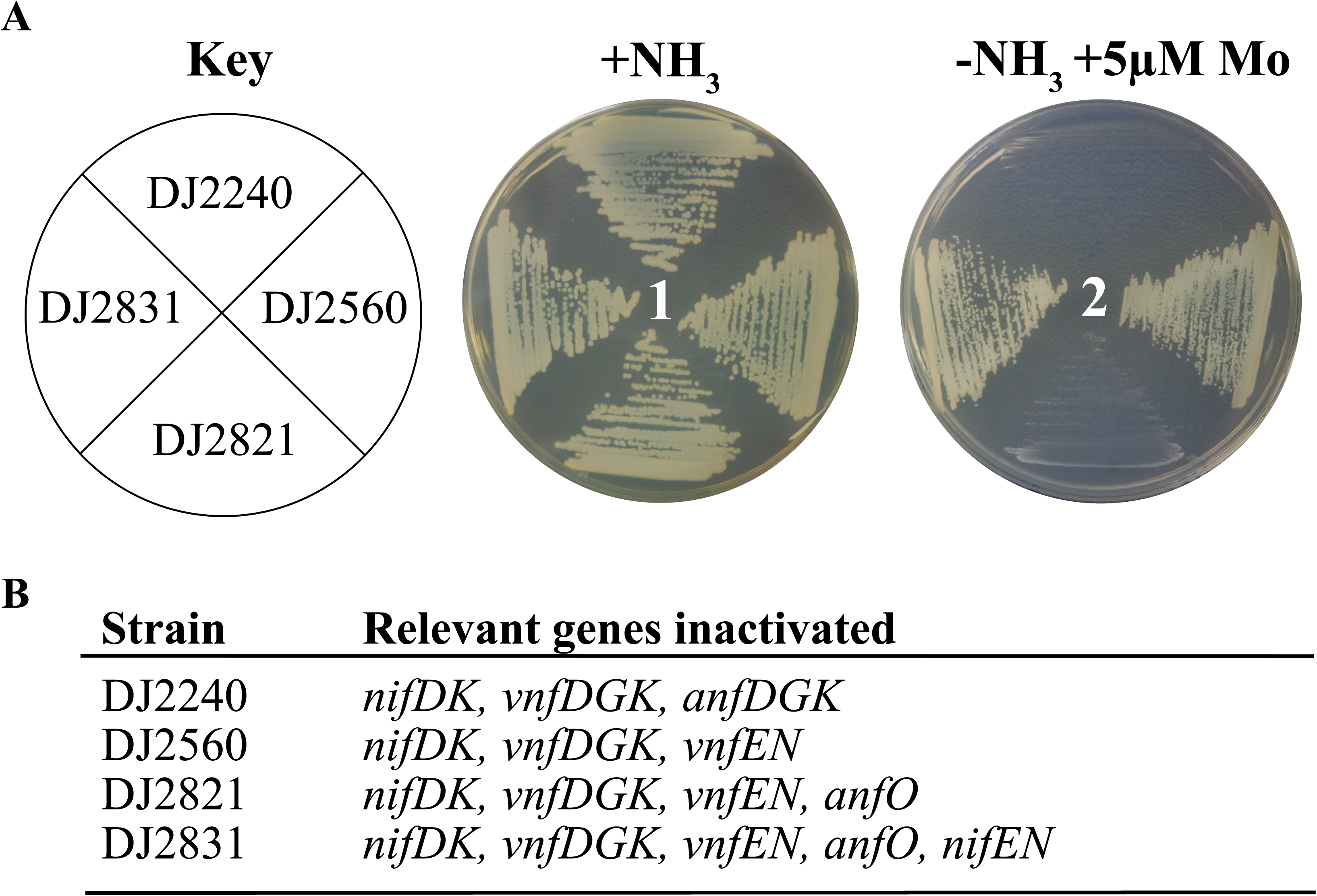
Phenotypic characterization of different *Azotobacter vinelandii* strains. A) Strains requiring Fe-only nitrogenase for diazotrophic growth were cultured on Burk’s medium agar plates containing a fixed nitrogen source (+NH_3_) (plate 1) or under diazotrophic conditions (−NH_3_) with Mo added to 5 μM (plate 2). B) Relevant genes inactivated for each strain. A complete genotypic description of each strain is indicated in Tables S1 and S2. The key observations are the inability of DJ2821 (Δ*anfO*) to grow under diazotrophic conditions when Mo is added to the growth medium and the reversal of that phenotype when both *anfO* and *nifEN* are inactivated (DJ2831), as shown in plate 2.

The catalytic properties of FeFe protein from three strains were evaluated (Table 1): a control strain (DJ2560) grown in the absence of Mo, a strain inactivated for *anfO* (DJ2821) grown in the presence of 1 μM Mo, and a strain inactivated for both *anfO* and *nifEN* (DJ2831) grown in the presence of 1 μM Mo. Cells were grown in large-scale cultures and FeFe protein was purified from cell paste as previously described (16). FeFe protein isolated from DJ2821(ϕ..*anfO*) cultured in the presence of 1μM Mo is unable to reduce N_2_. This observation is consistent with the inability of this strain to sustain diazotrophic growth when cultured in solid media in the presence of 5 µM Mo. It is also compatible with the possibility that DJ2821 produces a hybrid FeFe protein that contains FeMo-cofactor rather than FeFe-cofactor. The FeFe protein produced by DJ2821 grown in the presence of 1 µM Mo retains a capacity for proton and acetylene reduction, although these activities are lower when compared to the fully active FeFe protein isolated from DJ2560 (Table 1). It has been previously shown that FeFe protein having its normal complement of FeFe-cofactor has the capacity to reduce acetylene by both two and four electrons to yield ethylene and ethane, respectively (18, 20, 21). In contrast, MoFe protein containing its normal complement of FeMo-cofactor only catalyzes the two-electron reduction of acetylene to yield ethylene. Hybrid FeFe protein produced by DJ2821 containing FeMo-cofactor retains a capacity for two- and four-electron reduction of acetylene (Table 1). However, in the case of the hybrid FeFe-protein, the relative allocation of electrons to produce ethylene and ethane (∼1:1) is quite different than FeFe protein containing FeFe-cofactor (45:1). Thus, the protein environment as well as the identity of the heterometal within the catalytic cofactor (Fe or Mo) contribute to a capacity for the four-electron reduction of acetylene and the relative distribution of electrons to produce ethylene and ethane. Strikingly, nearly full FeFe protein directed N_2_ reduction, proton reduction, and acetylene reduction activities are recovered for FeFe protein from DJ2831, inactivated for both *anfO* and *nifEN*, even when grown in the presence of 1 µM Mo (Table 1). This result is consistent with reversal of the null diazotrophic growth phenotype when *nifEN*, required for FeMo-cofactor formation, is inactivated in combination with an *anfO* deletion (Fig. 3).

**Table 1.**
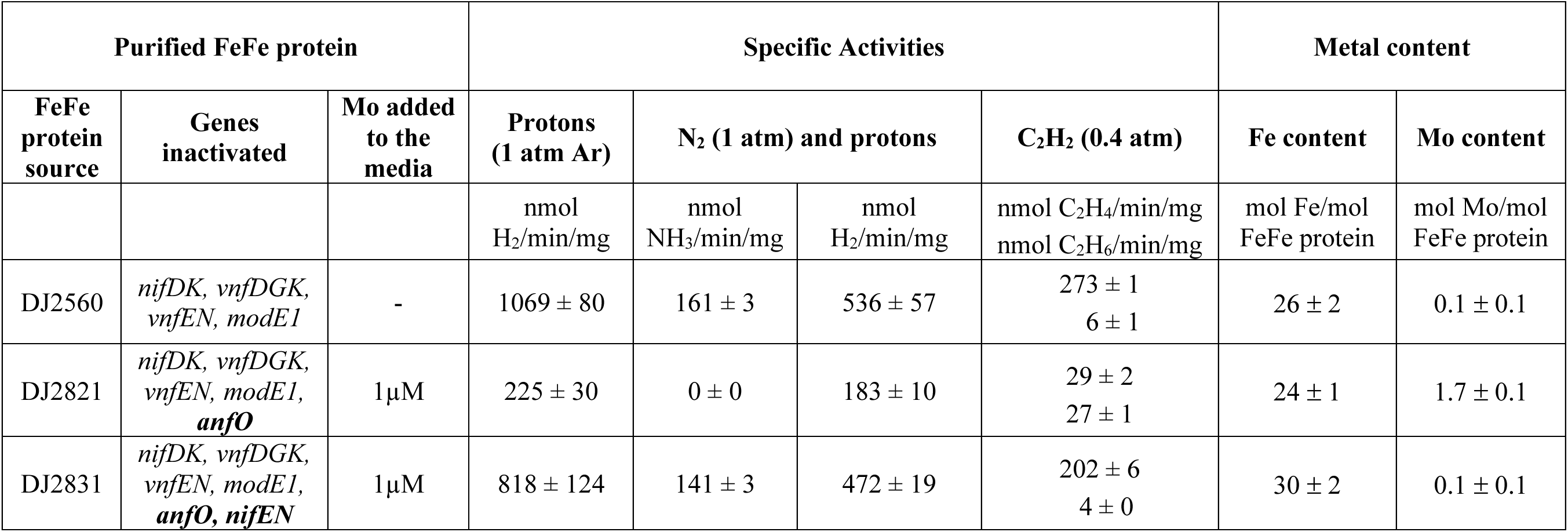
Catalytic properties and metal content of the FeFe proteins isolated from the different genetic backgrounds. FeFe proteins were purified from cells grown in the absence or presence of 1 μM Mo as indicated. Metal contents were quantified by ICP-MS. Molar ratios were calculated based on the molecular weight of the FeFe protein α_2_β_2_δ_2_ complex. Data presented are the average from at least three independent determinations. All strains contain genomic insertions or deletions in *nifDK*, *vnfDGK*, *vnfEN*, and *modE1* genes. Additional gene deletions are indicated in the table. See Tables S1 and S2 for detailed descriptions of the genotypes of DJ2560, DJ2821, and DJ2831.

| Purified FeFe protein |  |  | Specific Activities |  |  |  | Metal content |  |
| --- | --- | --- | --- | --- | --- | --- | --- | --- |
| FeFe protein source | Genes inactivated | Mo added to the media | Protons (1 atm Ar) | N<sub>2</sub> (1 atm) and protons |  | C<sub>2</sub>H<sub>2</sub> (0.4 atm) | Fe content | Mo content |
|  |  |  | nmol H<sub>2</sub>/min/mg | nmol NH<sub>3</sub>/min/mg | nmol H<sub>2</sub>/min/mg | nmol C<sub>2</sub>H<sub>4</sub>/min/mg<br>nmol C<sub>2</sub>H<sub>6</sub>/min/mg | mol Fe/mol FeFe protein | mol Mo/mol FeFe protein |
| DJ2560 | *nifDK*, *vnfDGK*, *vnfEN*, *modE1* | - | 1069 ± 80 | 161 ± 3 | 536 ± 57 | 273 ± 1<br>6 ± 1 | 26 ± 2 | 0.1 ± 0.1 |
| DJ2821 | *nifDK*, *vnfDGK*, *vnfEN*, *modE1*, *anfO* | 1 μM | 225 ± 30 | 0 ± 0 | 183 ± 10 | 29 ± 2<br>27 ± 1 | 24 ± 1 | 1.7 ± 0.1 |
| DJ2831 | *nifDK*, *vnfDGK*, *vnfEN*, *modE1*, *anfO*, *nifEN* | 1 μM | 818 ± 124 | 141 ± 3 | 472 ± 19 | 202 ± 6<br>4 ± 0 | 30 ± 2 | 0.1 ± 0.1 |

FeFe protein contains two complex metallocluster types: two 8Fe-7S P-clusters and two 8Fe-9S-C-homocitrate FeFe-cofactors (7, 21), giving a total of 32 associated Fe atoms in each fully loaded ɑ_2_β_2_δ_2_ FeFe protein heterohexamer (Fig. 1). FeFe protein that contains intact P-clusters, but lacks the catalytic FeFe-cofactor, does not contain the δ-subunit and is referred to as apo-FeFe protein (16). Loss of NifB-co formation results in accumulation of apo-FeFe protein. Metal analysis of FeFe protein isolated from the strains described above (Table 1) reveal that the Fe content of all three approach the theoretical values indicating they all contain P-clusters as well as a catalytic cofactor. However, for FeFe protein prepared from DJ2821 inactivated for *anfO* and cultured in the presence of 1 µM Mo there is also approximately 2 Mo for each FeFe protein hexamer (Table 1). This Mo content suggests FeFe protein produced by DJ2821 cultured in the presence of 1 µM Mo contains FeMo-cofactor rather than FeFe-cofactor. This possibility was confirmed by the observation that FeFe-protein prepared from DJ2831, inactivated for both *anfO* and *nifEN*, contains only very low levels of Mo as does FeFe protein produced by the control strain DJ2560 (Table 1).

The EPR spectra of FeFe protein isolated from each of the three strains (Fig. 4) confirm the hypothesis that the null diazotrophic activity of DJ2821 is due to misincorporation of FeMo-cofactor. In the presence of 2 mM dithionite, FeFe protein isolated from DJ2821 grown in the presence of 1 μM Mo prominently shows the characteristic *S* = 3/2 signal of FeMo-cofactor. Moreover, the *g*-values align well with an EPR spectrum from a previous study which assigned a similar signal to FeMo-cofactor misincorporation into FeFe-protein upon addition of molybdate to the growth media in the late-log and stationary phases of diazotrophically grown *Rhodobacter capsulatus* (21). Interestingly, the spectrum also shows a broad signal which could be due to improperly inserted FeMo-cofactor. The spectrum of FeFe protein isolated from DJ2560 grown without added Mo shows a small peak with the same *g*-values, indicating that this is also likely to result from misincorporated FeMo-cofactor generated from trace Mo in the media. FeFe protein from DJ2831 grown in the presence of 1 μM Mo shows no FeMo-cofactor signal, as expected due to the deletion of *nifEN*. By analogy to the V-dependent nitrogenase (22), the *g = 2* signals in these samples are likely to be associated with a P-cluster species. SDS-PAGE of the FeFe proteins purified from the different strains and growth conditions (DJ2560, DJ2821, and DJ2831) is shown in Fig. S1.

**Figure 4.**
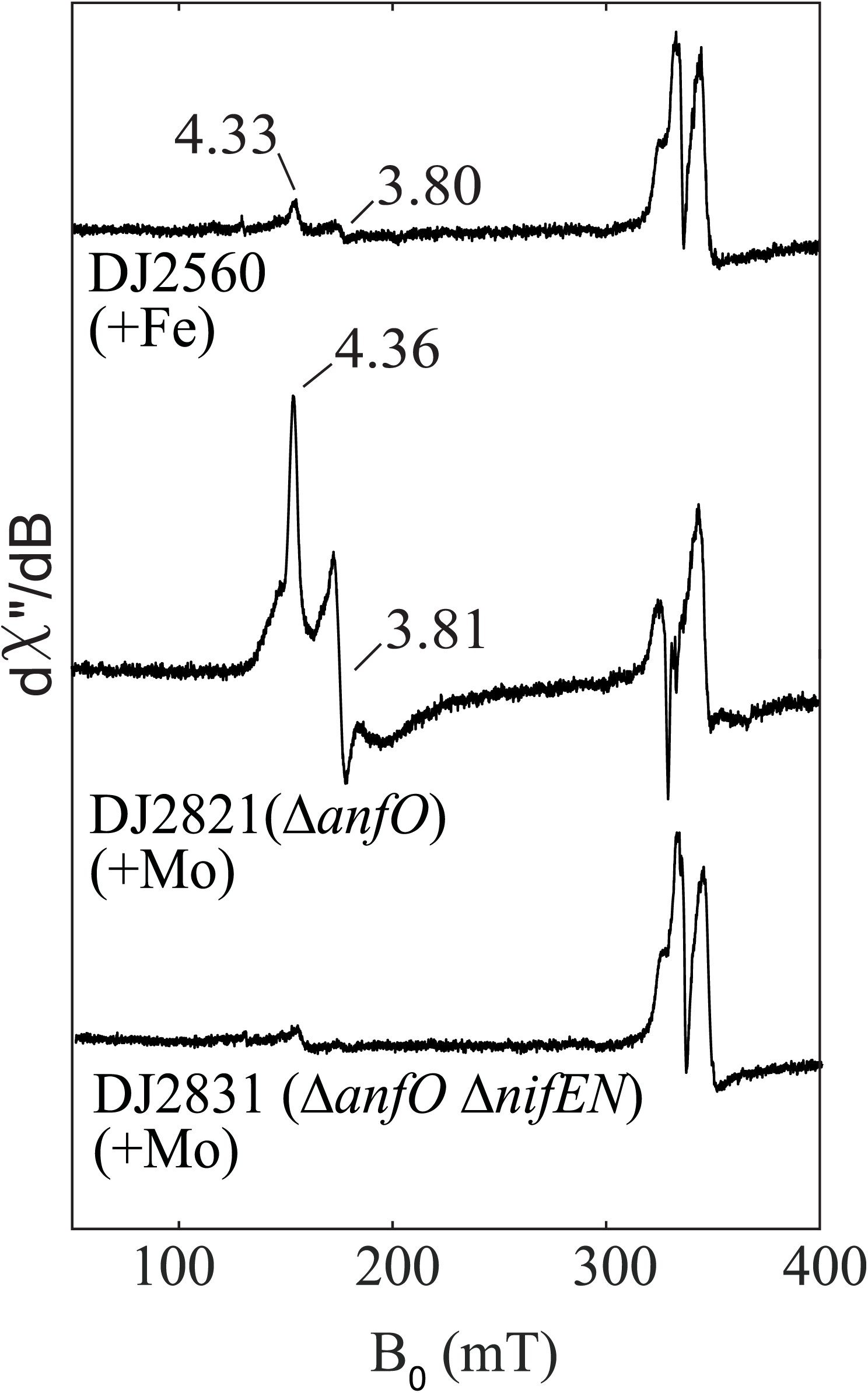
Comparison of the EPR spectra of isolated FeFe protein from *Azotobacter vinelandii* strains. Strep-tagged FeFe protein from strains DJ2560, DJ2821 (Δ*anfO*), and DJ2831 (Δ*anfO* Δ*nifEN*), cultured in the presence or absence of Mo, were purified and reduced with 2 mM dithionite. FeMo-cofactor in the reduced state exhibits an *S* = 3/2 signal, whereas FeFe-cofactor in the reduced state is EPR silent. FeFe protein purified from DJ2560 and DJ2821 exhibit FeMo-cofactor signals whose g-values match that of previously reported FeMo-cofactor in the FeFe protein of *Rhodobacter capsulatus* (19). The presence of FeMo-cofactor signal in DJ2560 is likely due to the presence of trace Mo contamination in the media as discussed in the results section. The EPR spectra were collected at 5 K, 9.37 GHz, and 63 μW. The *g* =2 signal present in all samples are associated with a P-cluster derived species.

Isolated FeFe protein containing FeFe-cofactor (obtained from DJ2241 grown in the absence of Mo and V), isolated FeFe protein containing FeV-cofactor (obtained from DJ2290 grown in the presence of 5 μM V (18), and FeFe protein containing FeMo-cofactor (obtained from DJ2821 grown in the presence of 1 μM Mo) are shown in Fig. S2. An equivalent amount of δ subunit is observed for all three isolated FeFe proteins indicating they are all replete with a catalytic cofactor, consistent with the requirement of catalytic cofactor insertion for δ subunit attachment (16).

### The N-terminal domain of AnfO binds apo-FeFe protein

Phylogenetic comparison of AnfO primary structures, as well as structural prediction based on Alphafold (23, 24), reveals that AnfO has two domains connected by a glycine-rich linker sequence indicating it could be a modular protein having two distinct functions (Fig. 5A). Comparative analysis of the predicted structure show that the AnfO N-terminal domain shares structural similarities with the predicted structures of other proteins having NifX-like domains, which are known or suspected to interact with immature nitrogenase isoforms, nitrogenase active-site cofactors, or their intermediates. These nitrogen fixation-associated proteins include NifX, VnfX, NifY, VnfY and NafY (25–30).

**Figure 5.**
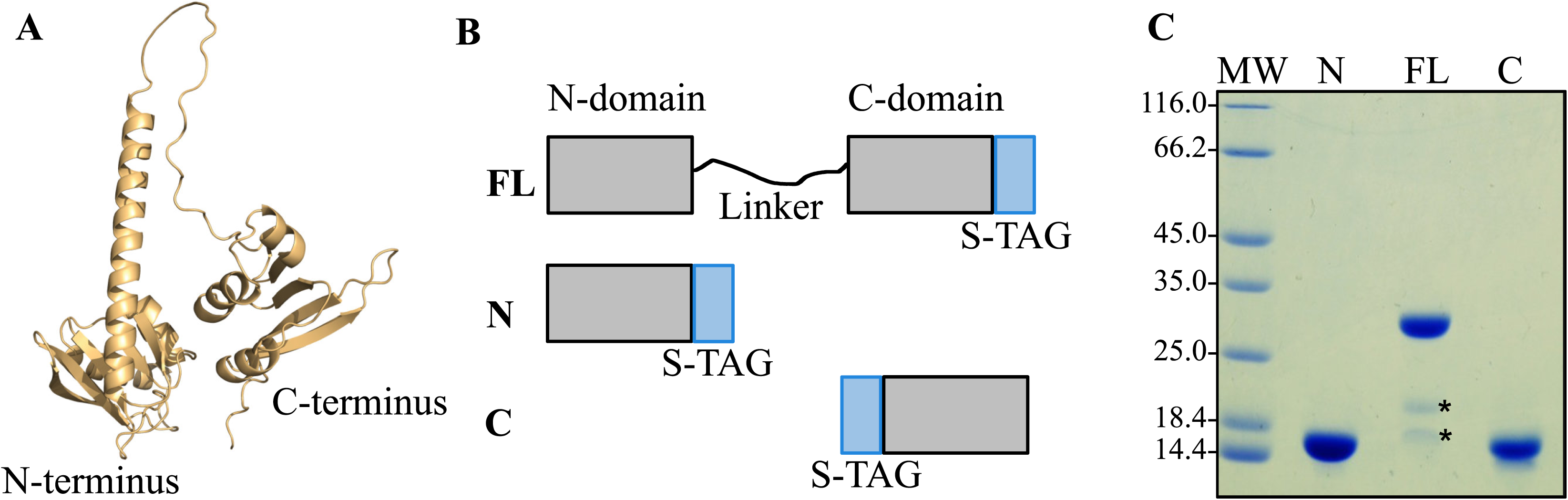
Structural features and isolation of heterologously produced AnfO and its two individual domains. A) Structural prediction of AnfO using Alphafold indicates it has two-domains connected by a linker region. B) Schematic representation of Strep-tagged (“S-TAG”) versions of AnfO heterologously produced in *E. coli* (“FL”: full-length, “N”: N-domain including AnfO residues 1-132, “C”: C-domain including residues 138-235). C) SDS-PAGE of the different versions of AnfO (N, FL, C) purified from *E. coli*. The two smaller proteins indicated by asterisks in the lane labeled “FL” were identified by Mass Spectrometry to be full-length AnfO cleavage products. The structure of AnfO was predicted using Colabfold using AlphaFold2 (61). Colabfold was run with the default 3 recycles. The three-dimensional model was visualized using PyMOL (58).

Genetic constructs revealed that placement of a Strep-tag at the C-terminus of AnfO (DJ2494) does not impair *in vivo* function whereas placement of a Strep-tag at the N-terminus (DJ2527) inactivates *in vivo* function based on the diazotrophic growth phenotypes of the corresponding strains (Fig. S3). Construction of a strain carrying an in-frame deletion within the C-terminal coding portion of *anfO* (DJ2821) also results in a null diazotrophic growth phenotype when cultured in the presence of 5 µM Mo (Fig. 3). Furthermore, substitution of the conserved AnfO His^203^ residue, a potential FeMo-cofactor/FeV-cofactor ligand, inactivates AnfO (Fig. S4). Thus, both the N-terminal and C-terminal domains are required to sustain AnfO function. Because the AnfO C-terminus can be modified without affecting *in vivo* function, a plasmid construct was used to place a Strep-tag at the AnfO C-terminus to enable its heterologous production and purification from *Escherichia coli*. AnfO produced and purified from this construct is designated “FL” to indicate full-length AnfO (Fig 5B). Two other truncated and Strep-tagged versions of AnfO were also produced and purified. One of these, designated “N-domain”, contains a Strep-tagged version of an AnfO fragment that includes amino acid residues 1-132 encompassing the N-terminal domain. The other, designated “C-domain”, contains a Twin-Strep-tagged version of an AnfO fragment that includes amino acid residues 138-245 (Fig. 5B). These three AnfO-derived species were separately expressed in and purified from *E. coli* cells (Fig. 5C) and subsequently used in bait-prey experiments to explore their possible interaction with FeFe protein. Cell extracts prepared from strain DJ2239, deleted for *nifDK* and *vnfDGK* and having no affinity purification tag placed on the FeFe protein, were passed over Strep-Tactin columns that separately contain immobilized FL, N- or C-domains followed by elution of the immobilized proteins (Fig. 6). The N-domain and FL protein both captured the FeFe protein α- and β-subunits, but not the δ subunit. As previously noted for both VFe protein and FeFe protein, NifB-co formation is required for the attachment of their corresponding δ subunits. The absence of the δ subunit therefore indicates the species captured by either AnfO or its N-terminal domain is apo-FeFe protein. No protein was captured by the C-domain.

**Figure 6.**
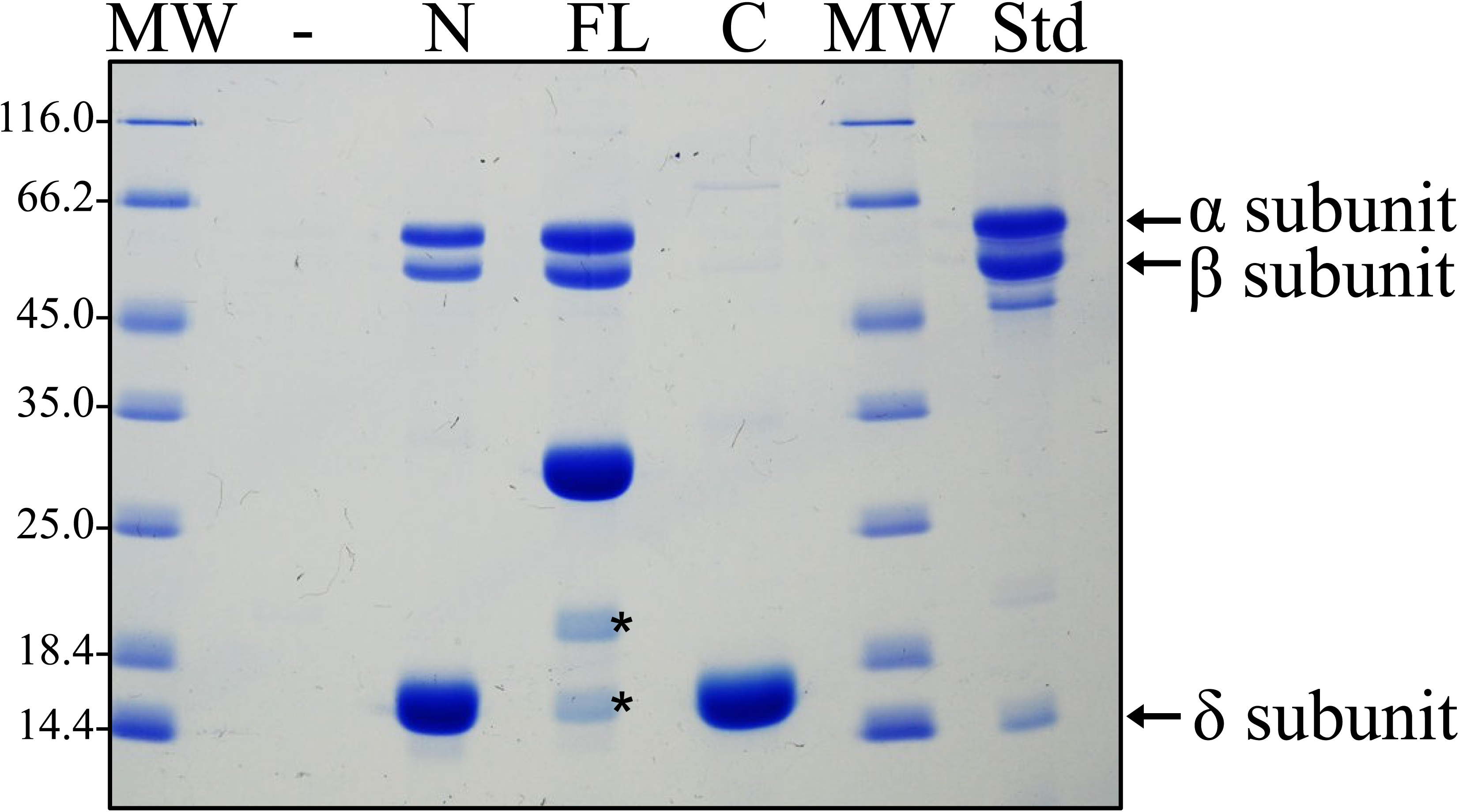
The N-terminal domain of AnfO interacts with apo-FeFe protein. Strep-tagged versions of purified full-length AnfO, its N-terminal domain, or its C-terminal domain produced in *E. coli* were used to saturate individual Strep-Tactin columns and washed with buffer B described in Experimental Procedures. Cell extract of DJ2239, which produces Fe-only nitrogenase but not the Mo-dependent or V-dependent isoforms, was then passed over each column and washed with buffer B containing no biotin, followed by elution with buffer B containing 50 mM biotin. Coomassie blue stained SDS-PAGE analysis of the eluted samples is shown. Full-length AnfO and its N-domain capture the FeFe protein α- and β-subunits but not the FeFe protein 8 subunit, whereas the AnfO C-domain did not capture any other proteins. The second lane represents a negative control for which no bait protein was immobilized on the Strep-Tactin column. MW = molecular weight standards (kDa), FL = immobilized full-length AnfO used as bait, N = immobilized AnfO N-domain used as bait, C = immobilized AnfO C-domain used as bait, Std = purified FeFe protein standard with the αβ8 subunits indicated by arrows. The two smaller proteins indicated by asterisks in the lane labelled “FL” are full-length AnfO cleavage products. The identity of all proteins was confirmed by Mass Spectrometry.

To further confirm that AnfO binds apo-FeFe protein, another bait-prey experiment was performed following the methodology already described. In this case, AnfO FL, N-, or C-domains were immobilized on separate Strep-Tactin columns and crude extract containing apo-FeFe protein prepared from a strain deleted for *nifB* was passed over the corresponding columns. Apo-FeFe protein, which does not contain the δ-subunit, was captured by FL AnfO and its N-terminal domain but was not captured by the C-terminal domain (Fig. S5). A complementary experiment using immobilized Strep-tagged apo-FeFe protein produced by a NifB-deficient strain as bait was also performed. Immobilized apo-FeFe protein was able to capture FL AnfO from crude extracts of AnfO recombinantly produced in *E. coli* (Fig. S6).

### AnfO C-terminal domain can bind FeMo- and FeV-cofactor

The experiments described above demonstrate that the N-terminal domain of AnfO is responsible for binding apo-FeFe protein. We next performed experiments to shed light on how AnfO prevents catalytic cofactor misincorporation, with the specific goal of understanding if AnfO binds FeMo-cofactor and/or FeV-cofactor, and if so, which domain of AnfO is responsible for cofactor binding. At the foundation of these experiments are previous studies that have demonstrated that FeMo-cofactor and FeV-cofactor can be isolated in organic solvents (specifically, *N*-methylformamide (NMF)) and re-inserted into various proteins (31–33). Here, we conducted similar studies using AnfO and its truncated forms as the protein host.

FeMo-cofactor was isolated from MoFe protein following reported protocols (31, 34, 35) and subsequently incubated with 1.1 equivalent of FL AnfO. The resulting FL AnfO protein was purified from any small molecules by concentrating the mixture >1000-fold to yield a brown sample, indicating that the protein now harbored an Fe-S cluster. ICP-MS analysis revealed an Fe:Mo ratio of 7.1:1 (which compares favorably to the theoretical value of 7:1 for FeMo-cofactor) and a 0.51:1 FeMo-cofactor:polypeptide ratio (Table S4). Furthermore, EPR spectroscopic analysis showed an *S* = 3/2 signal that is similar to, but clearly distinct from, that of isolated FeMo-cofactor (Fig. 7B). Taken together, these findings demonstrate that FL AnfO binds FeMo-cofactor. Similar experiments were conducted using FeV-cofactor, which was isolated from VFe protein following reported protocols (33). FeV-cofactor loaded FL AnfO also showed an intense brown color after >1000-fold concentration. ICP-MS analysis revealed an Fe:V ratio of 8.6:1, which compares favorably to the theoretical 7:1 Fe:V ratio of FeV-cofactor. The EPR spectrum showed an *S* = 3/2 signal similar to isolated FeV-cofactor, demonstrating that FL AnfO indeed is capable of binding FeV-cofactor (Fig. 7C).

**Figure 7.**
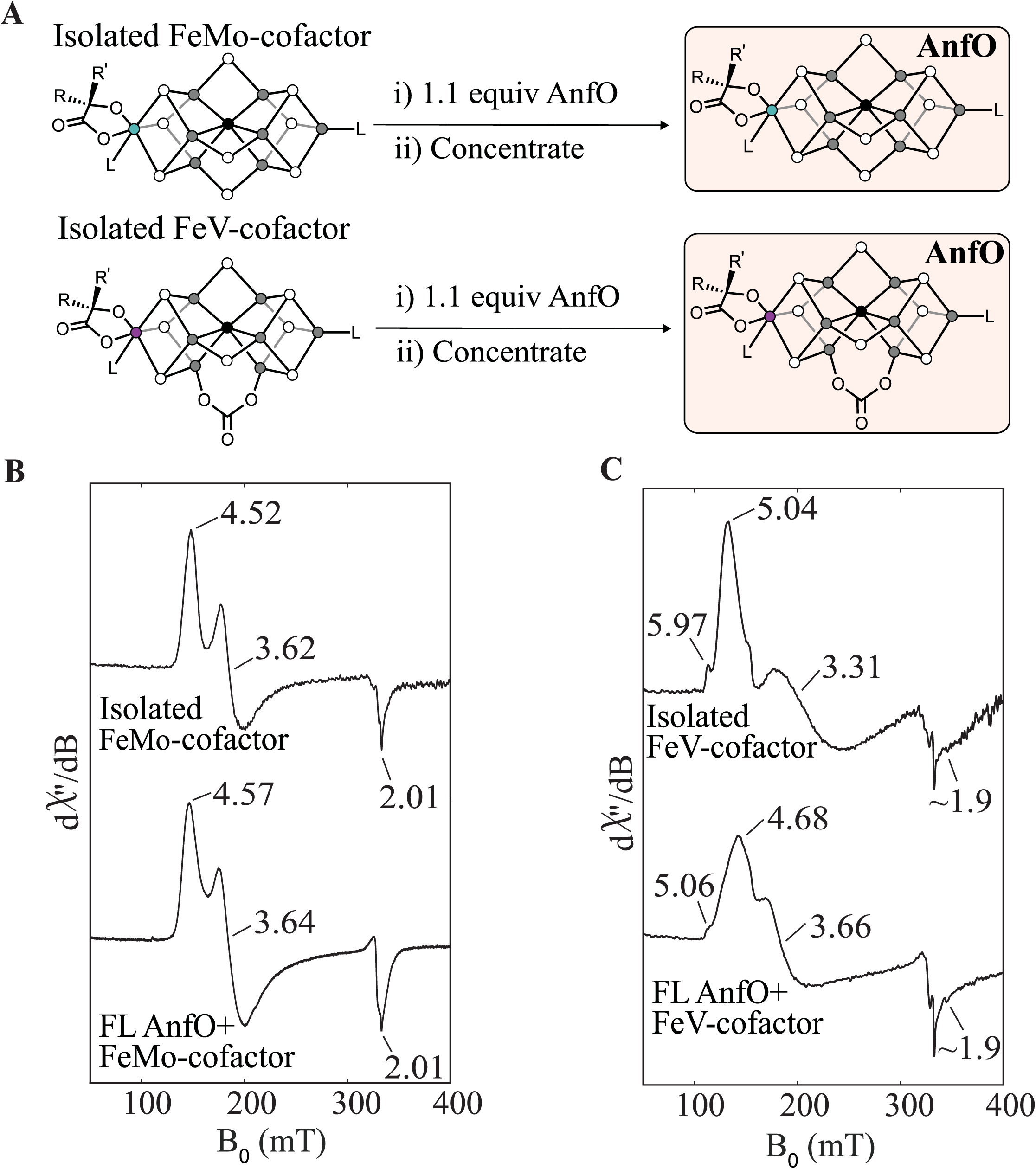
Cofactor binding to full length AnfO. A) Schematic representation of the FeMo- or FeV-cofactor binding to full length AnfO. B) EPR spectra of FeMo-cofactor (in NMF with 1 mM PhSH) and of full length AnfO after incubation with FeMo-cofactor and repurification. See Methods for details. C) EPR spectra of FeV-cofactor (in NMF with 1 mM PhSH) and of full length AnfO after incubation with FeV-cofactor and repurification. See Methods for details.

Further evidence for the specific binding of nitrogenase cofactors by FL AnfO was provided by comparison of the EPR spectra of FeMo-cofactor bound to FL AnfO, NifX, and NafY (Fig. 8). NafY is involved in intracellular trafficking of FeMo-cofactor and is known to bind FeMo-cofactor (17). Although NifX is similarly involved in the intracellular trafficking of NifB-co (17), it also has the *in vitro* capacity to bind FeMo-cofactor as shown here. Each spectrum features an *S* = 3/2 signal that has slightly different effective *g*-values (*g*_eff_ values), representing different ratios and distributions of the zero-field splitting parameters, *D* and *E*. The different *E*/*D* ratios and lineshapes of each signal result from differences in the local environments of FeMo-cofactor and, collectively, indicate that FeMo-cofactor binds in a manner that is specific to each polypeptide. We therefore rule out non-specific binding of FeMo-cofactor to AnfO based on the well-defined line shapes of the EPR spectra and unique *g*-values, and conclude that one function of FL AnfO is to bind FeMo-cofactor and FeV-cofactor.

**Figure 8.**
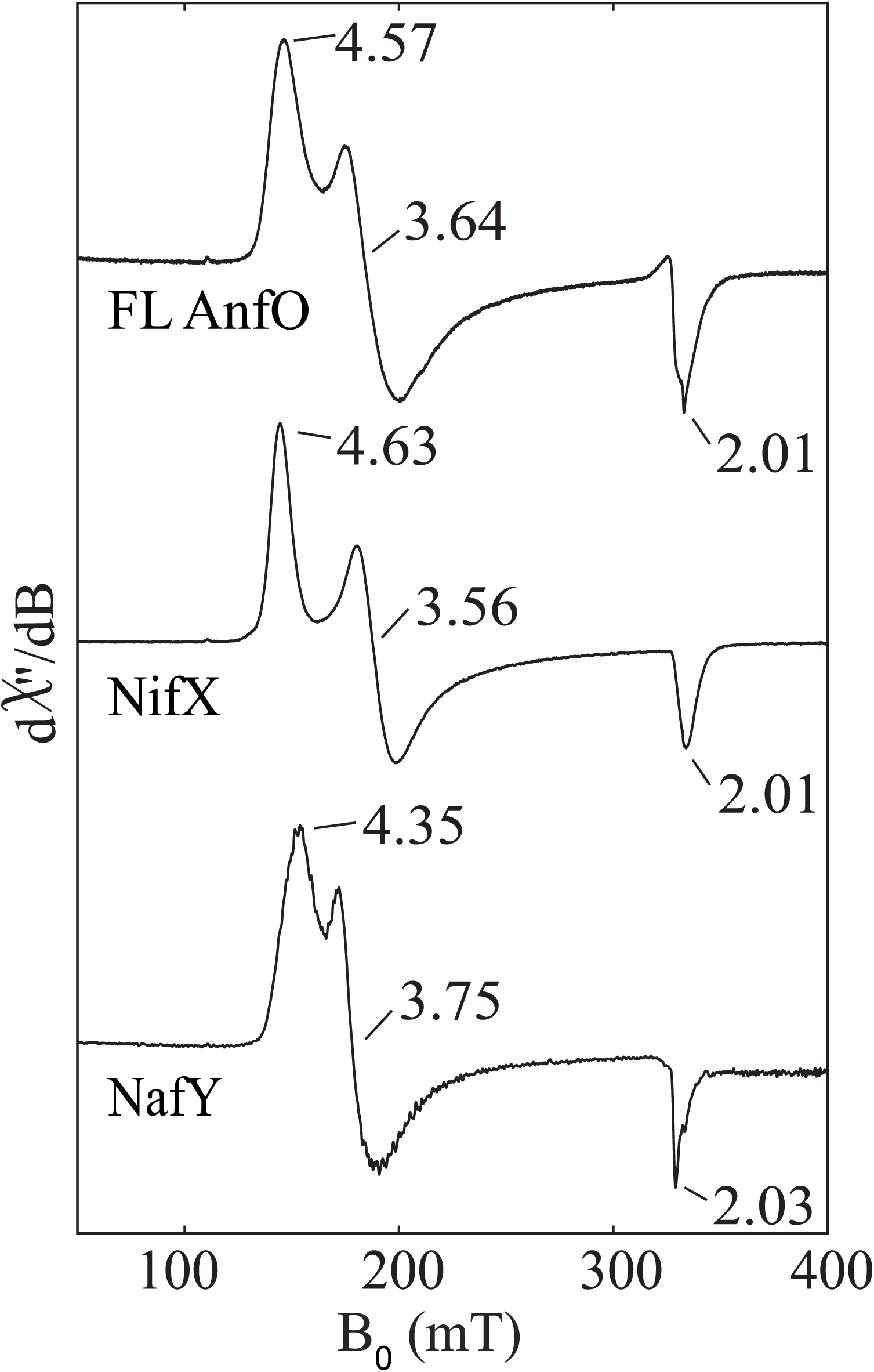
Comparison of the EPR spectra of FeMo-cofactor bound to full length AnfO, NifX, and NafY. “FL AnfO”: full length AnfO. EPR conditions: 9.37 GHz, 5 K, 1 mW. See Methods for details.

Given the above findings with FL AnfO, we next sought to determine which domain is responsible for catalytic cofactor binding. The individual affinity-tagged N- and C-domains were incubated with isolated FeMo-cofactor and applied to separate Strep-Tactin affinity columns. A control sample was also prepared in which isolated FeMo-cofactor was applied to a Strep-Tactin affinity column. For this sample it was found that free FeMo-cofactor becomes irreversibly bound to the column and cannot be eluted. Consequently, for experimental samples, any FeMo-cofactor co-eluting with bound proteins must be associated with that protein species because any free FeMo-cofactor will remain bound to the column. Fig. 9A shows that, after column washing and prior to elution using a biotin-containing buffer to free protein samples from the affinity resin, the top of each column exhibits a brown color. After elution, the characteristic brown color associated with FeMo-cofactor in the control sample remained at the top of the Strep-Tactin affinity column, as already noted. In the case of the N-domain sample, FeMo-cofactor also remained immobilized at the top of the column following elution indicating that no FeMo-cofactor was captured by the N-fragment. Furthermore, the N-domain eluate was nearly colorless also indicating no FeMo-cofactor co-elutes with this species. In contrast, FeMo-cofactor does co-elute with the C-domain as shown by loss of the brown color from the top of the column upon elution and appearance of a brown eluate. These observations indicate that the C-terminal domain of AnfO, and not the N-terminal domain, is responsible for binding FeMo-cofactor. Further support for this conclusion was obtained by incubating the C-domain with isolated FeMo-cofactor and analyzing the purified protein. The observed Fe:Mo ratio of the species captured by the AnfO C-domain, 7.1:1, is close to the theoretical 7:1 ratio for FeMo-cofactor. The EPR spectrum also reveals an *S* = 3/2 signal similar to that of FeMo-cofactor loaded FL AnfO (Fig. 9B).

**Figure 9.**
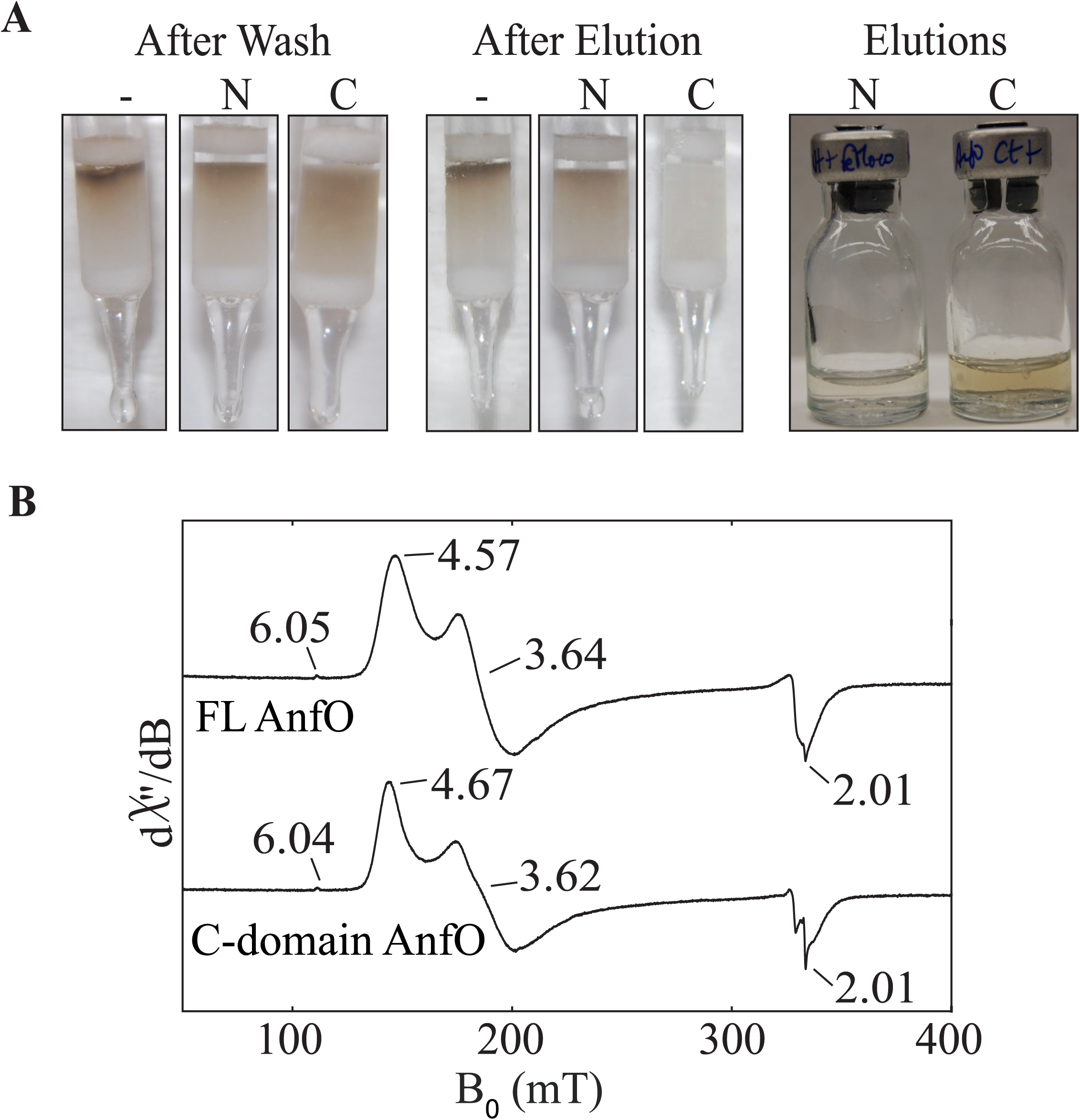
The C-terminal domain of AnfO interacts with isolated FeMo-cofactor. Prior to their application to a Strep-Tactin affinity column, individually isolated N-domain and C-domain proteins were mixed with FeMo-cofactor as described in Experimental Procedures. Samples were then applied to Strep-Tactin columns and washed with several column volumes of Buffer B (described in Experimental Procedures). Samples were subsequently eluted with Buffer B containing 50 mM biotin. Isolated FeMo-cofactor was also applied to a Strep-Tactin column to serve as a control. The leftward Panel in (A) shows the appearance of the columns after washing with Buffer B, the central Panel in (A) shows the appearance of the columns after elution with Buffer B containing 50 mM biotin, and the rightward panel in (A) shows the appearance of the eluates. The control sample is indicated by (-), and the samples having either the AnfO N-domain or C-domain mixed with isolated FeMo-cofactor are indicated as (N) and (C), respectively. Note that in the control sample FeMo-cofactor becomes irreversibly bound to the Strep-Tactin column. FeMo-cofactor is also retained on the column for the sample containing the mixture of FeMo-cofactor and AnfO N-domain, indicating the FeMo-cofactor does not bind to the AnfO N-domain. The brown color evident in the AnfO C-domain eluate in the rightward Panel indicates binding of FeMo-cofactor to AnfO. B) EPR spectra (9.37 GHz, 5 K, 1 mW) of full length AnfO (“FL”) and C-domain AnfO bound to FeMo-cofactor showing that the two spectra are nearly identical.

To explore ligands that could be involved in the capture of FeMo-cofactor and FeV-cofactor by AnfO, conserved residues Cys^159^, Cys^201^, and His^203^ were respectively substituted by Ala^159^, Ala^201^ and Leu^203^. This analysis revealed that substitution of His^203^ inactivated the *in vivo* function of AnfO whereas substitution of neither Cys^159^ nor Cys^201^ altered the *in vivo* capacity for AnfO to preserve the fidelity of FeFe protein maturation (Fig S6). These results are similar to amino acid substitutions involving the NifB-co trafficking protein NifX for which the conserved His^35^ residue, but not the conserved Cys^82^ residue, is involved in binding NifB-co (35). They also resemble the situation of the amino acid substitutions of NafY, for which the His^121^ residue is the essential to bind FeMo-cofactor (32, 36).

## Discussion

The structural similarity of the three nitrogenase catalytic cofactors presents a substantial challenge to the cell, especially during the transition to utilization of the Fe-only isoform: namely, how can the misincorporation of FeMo-cofactor or FeV-cofactor into apo-FeFe protein be prevented. Processing of the universal precursor, NifB-co, to form FeMo-cofactor and FeV-cofactor occurs on scaffolds, followed by their insertion into apo-forms of their cognate protein partners. In the case of FeFe-cofactor, which only requires the attachment of homocitrate to NifB-co, no scaffold is involved, suggesting homocitrate attachment occurs directly within an immature form of the FeFe protein. Under steady-state growth conditions, differential accumulation of the nitrogenase isoforms is exquisitely controlled at the transcriptional level in response to a physiological demand for nitrogen fixation and metal availability (17). However, because FeMo-cofactor and FeV-cofactor are separately synthesized and then inserted into apo-forms of their cognate proteins, it is likely that a pool of pre-formed FeMo-cofactor or FeV-cofactor cofactor is available under growth conditions that favor accumulation of the Mo-dependent or V-dependent nitrogenase isoforms. Once neither Mo nor V is available, cells must transition to utilization of the Fe-only isoform. During the initial stages of such a transition, the pool of preformed FeMo-cofactor or FeV-cofactor would be available to become inserted into an immature apo-form of FeFe protein, rendering the protein inactive for N_2_ reduction, and thereby causing a delay in adjustment to the emergent growth condition. The present work shows that an auxiliary protein, AnfO, which accumulates under conditions that favor Fe-only nitrogenase utilization, preserves the fidelity of FeFe protein maturation by preventing such misincorporation. AnfO might also serve this role in situations where more than one isoform is co-expressed at the same time. For example, Mo- and Fe-only nitrogenases are co-expressed under low Mo conditions in *Rhodobacter capsulatus* (21, 37, 38), while it has been reported that *Methanosarcina acetivorans* (39) is able to produce the three nitrogenase isoforms simultaneously when grown in Mo-depleted conditions. *R. capsulatus* and *M. acetivorans* both express *anfO* in association with Fe-only nitrogenase (40). Given their nearly identical structures, charge, and topologies it is highly unlikely that AnfO could preferentially bind FeMo-cofactor and FeV-cofactor but not FeFe cofactor. Instead, why AnfO can prevent misincorporation of FeMo-cofactor and FeV-cofactor into immature FeFe protein while not affecting the normal maturation process to yield FeFe protein having FeFe-cofactor is likely associated with the difference in the assembly pathways for the corresponding cofactors (Fig. 2). Namely, because FeFe-cofactor assembly is completed directly within an immature form of FeFe protein there is never free FeFe-cofactor available for capturing by AnfO. Thus, both differences in corresponding catalytic cofactor assembly processes and the ability of AnfO to bind free FeMo- and FeV-cofactors to prevent their misincorporation into immature FeFe protein represent complementary strategies to assure the fidelity of FeFe protein maturation.

Structural considerations and biochemical characterization of AnfO provide mechanistic insights into how it preserves the fidelity of FeFe protein maturation. AnfO contains two domains connected by a glycine-rich flexible linker region: the N-terminal domain which has the capacity to bind immature cofactor-less FeFe protein, and the C-terminal domain which can bind FeMo-cofactor and FeV-cofactor. These observations lead to a model during FeFe protein maturation wherein the N-terminal domain tethers AnfO to immature FeFe protein, thereby effectively localizing the C-terminal domain to capture FeMo-cofactor and FeV-cofactor to prevent their misincorporation into the immature FeFe protein (Fig. 10). Localization of the C-terminal domain, accomplished by tethering of the N-terminal domain to immature FeFe protein, is an important aspect of protection because both domains are required *in vivo* to preserve the fidelity of FeFe protein maturation. Dissociation of AnfO is predicted to occur upon insertion of NifB-co or completion of FeFe-cofactor formation by homocitrate insertion followed by attachment of the δ-subunit. The attachment of the δ-subunit is already known to require NifB-co availability (16). A plausible description of these events is shown in Figure 10. This situation resembles the function of the two-domain protein, NafY, in the Mo-nitrogenase system. In this case, NafY interacts with FeMo-cofactor and with apo-MoFe protein (25, 32, 36) to facilitate delivery of FeMo-cofactor from the NifEN assembly scaffold to apo-MoFe protein (36), with the attendant dissociation of NafY upon FeMo-cofactor insertion. It is likely that a protein having sequence similarity to NafY, designated VnfY, has the same function as NafY involving the trafficking of FeV-cofactor for maturation of VFe protein (29).

**Figure 10.**
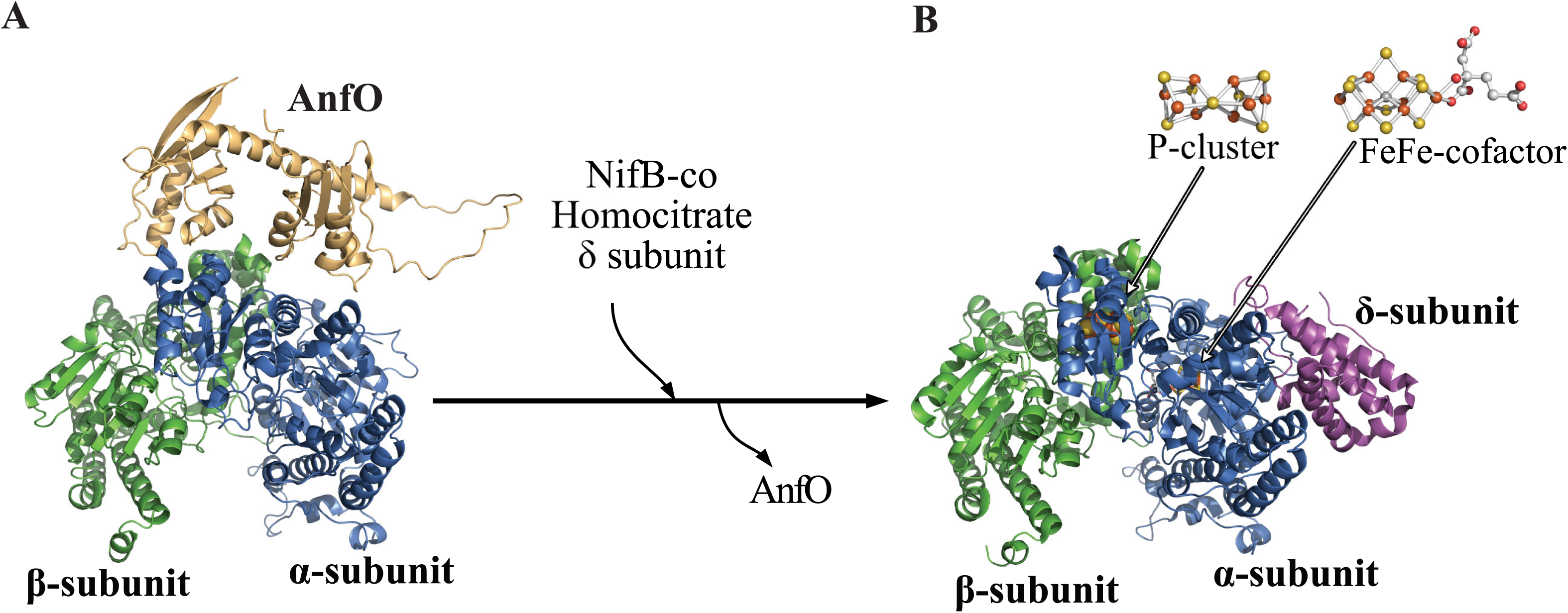
Conceptual framework for protection of immature FeFe protein from misincorporation of FeMo-cofactor and FeV-cofactor. A) The N-terminal domain of AnfO interacts with apo-FeFe protein while the C-terminal domain, tethered by a flexible linker is poised to capture FeMo- or FeV-cofactor to prevent their misincorporation. B) Incorporation of NifB-co, or completion of FeFe-cofactor formation by attachment of homocitrate within FeFe protein (see Fig. 2), results in the dissociation of AnfO and the attachment of the δ-subunit to the apo-FeFe protein. Structures are represented as follows: blue: FeFe protein α-subunit; green: FeFe protein β-subunit; pink: FeFe protein δ-subunit; light orange: AnfO. For convenience, only one half of the FeFe protein is represented. In (B) the P-cluster is shown at the αβ interface and FeFe-cofactor is located within the α-subunit. The holo-FeFe protein structure was extracted from PDB 8BOQ (7). The apo-FeFe protein structure was predicted using ColabFold (61). The AnfO structure and its favored interaction with apo-FeFe protein was predicted using Alphafold (23, 24). The P-cluster was not modeled into the apo-FeFe protein interacting with AnfO shown in Panel A. The three-dimensional models were visualized using PyMol (58).

The present work also provides a fresh opportunity to explore the contribution of the heterometal site contained in the nitrogenase catalytic cofactors to catalysis. Previous work established that hybrid species of various nitrogenase isoforms containing the incorrect or incomplete catalytic cofactor generally preserve a capacity for proton and acetylene reduction but are inactive for N_2_ reduction (18, 41–45). This alteration in reactivities is also the case for hybrid FeFe protein containing FeMo-cofactor, which supports proton and acetylene reduction at modest levels but cannot reduce N_2_. This is a remarkable observation given that the chemical compositions of FeFe-cofactor and FeMo-cofactor are identical with the exception that the apical metal atom attached to homocitrate is either Fe or Mo. Moreover, the specific contribution of the apical heterometal contained in the various cofactors to catalysis remains unknown. Because the accumulation of more electrons is required for N_2_ activation compared to proton or acetylene reduction (46), the catalytic features of the hybrid FeFe protein described here suggest that the apical metal species plays an important role in achieving or stabilizing the state required for N_2_ activation. Whether or not this feature is the consequence of subtle differences in the geometric configurations of FeMo-cofactor and FeFe-cofactor when contained in the FeFe protein chassis, or different electronic features imposed by the nature of the heterometal, remains to be explored. The ability to produce a hybrid FeFe protein containing FeMo-cofactor at scale, together with knowledge of the three-dimensional structures of all three nitrogenase isoforms (47), now provides a rich source for the biophysical and enzymological elucidation of the role of the apical metal atom contained in the catalytic cofactors.

## Experimental procedures

### Strains and plasmids used in this work

*A. vinelandii* strains used in this work are listed in Table S1. All strains are tungsten tolerant. For strains derived from DJ1254 W tolerance is the result of a 42,096-bp genomic deletion (48, 49), and for strains derived from DJ2560 W tolerance is the result of a *modE1* deletion (16). For all strains a Strep-tag encoding DNA sequence giving the peptide sequence ASWSHPQFEK was placed at the C-terminus of *anfD* for affinity purification purposes. The precise location of insertions and/or deletions for each strain are listed in Table S2, which also indicates the plasmids used for the construction of the corresponding strains. For convenience, deletion/insertion nomenclature for each plasmid corresponds to the genotypic description shown in Table S1. *A. vinelandii* transformations were performed as previously described (50). Genomic insertions/deletions were confirmed by PCR amplification and/or sequence determination of the appropriate genomic region. DNA sequence analysis was performed by the Genomics Sequencing Center service provided by the Fralin Life Sciences Institute, Virginia Tech.

For heterologous expression of AnfO, AnfO N-domain, and AnfO C-domain, *E. coli* BL21(DE3) cells were transformed with pT7-7 derived recombinant plasmids shown in Table S3. A DNA sequence giving the peptide sequence ASWSHPQFEK (Strep-tag) or ASWSHPQFEKGGGSGGGSGGSAWSHPQFEKAS (Twin Strep-tag) was placed at the N- or C-terminus of these constructs as indicated in the appropriate figure legends.

### Bacterial growth

*A. vinelandii* cells were cultured at 30°C on Burk’s modified agar plates (51) with or without supplementation of 13 mM ammonium acetate (Sigma-Aldrich), or 5 µM Na_2_MoO_4_ (J.T. Baker, Phillipsburg, NJ) as indicated in the tables and figures. Bacto Agar (BD, Sparks, MD), was used as the solidifying agent. For large-scale liquid cultures, *A. vinelandii* cells were grown in a 150L custom-built fermenter (W.B. Moore, Inc., Easton, PA) at 30 °C in a modified Burk medium containing 1mM urea as the nitrogen source. Where indicated, 1 µM Na_2_MoO_4_ was included in the media. Cells were grown overnight and harvested at an OD_600_ between 1.2 and 2.

*E. coli* BL21(DE3) chemically competent cells were transformed using recombinant plasmids listed in Table S3 and cultured at 37 °C in LB solid media plates supplemented with ampicillin (100 µg/ml). For liquid cultures, *E. coli* cells harboring the appropriate heterologous expression plasmid were grown in 6 L of LB media supplemented with ampicillin and grown at 37 °C. When cultures reached OD_600_ ∼ 0.6 lactose was added to give a final concentration of 10 g/L to induce heterologous expression of AnfO or an AnfO domain. Following lactose induction, cells were incubated at 30 °C for an additional 3 h and harvested.

### Protein purification and analysis

FeFe proteins produced in *A. vinelandii* were purified following previously described procedures using Strep-Tactin columns (IBA Lifesciences, Gottingen, Germany) (18, 52). Strep-tagged AnfO or AnfO domains produced in *E. coli* were initially isolated using a previously described affinity purification procedure (52) and further purified using a HiTrap Q-sepharose or HiTrap desalting column (Cytiva, Uppsala, Sweden) to remove biotin. Protein concentrations were determined by the BCA method (BCA protein assay kit, Sigma-Aldrich, Saint Louis, MO). Protein samples that contained dithionite (Na_2_S_2_O_4_) in the elution buffer were incubated with iodoacetamide (8 mg/mL) for at least 30 minutes at 37 °C prior performing the BCA assay (53). Protein purity was evaluated by Coomassie blue stained SDS-PAGE analysis. Molecular weight standards: β-galactosidase (116.0 kDa), Bovine serum albumin (66.2 kDa), Ovalbumin (45 kDa), Lactate dehydrogenase (35 kDa), REase Bsp981 (25 kDa), β-lactoglobulin (18.4 kDa) and Lysozyme (14.4 kDa) were obtained from Fisher Scientific (Waltham, MA). Metal content (Fe, Mo, and V) was determined by inductively coupled plasma mass spectrometry (ICP-MS) operated by the MIT or the Virginia Tech Metal Analysis Service.

### Substrate reduction assays

Substrate reduction assays were performed using sealed 9.4 mL serum vials containing 1 mL assay cocktail, and purified FeFe protein and Fe-protein 3, as previously described (54). Headspace gasses in the vials were adjusted to the desired partial pressures of a particular gaseous substrate (N_2_, C_2_H_2_, as indicated) and remaining space was filled with Ar. Each 1 mL assay cocktail contained 6.7 mM MgCl_2_, 30 mM phosphocreatine, 5 mM ATP, 0.2 mg/mL creatine phosphokinase, 10 mM dithionite, 100 mM MOPS (pH 7.0) and 0.9 mg Fe protein-3 (AnfH). Reactions were initiated by the addition of 0.1 mg FeFe protein. Vials were then ventilated to atmospheric pressure and incubated with agitation at 30°C for 8 min, and quenched by the addition of 300 μL of 0.4M EDTA (pH 8.0) to stop the reaction. Substrate reduction products NH_3_, H_2_, C_2_H_4,_ and C_2_H_6_ were quantified as described previously (55–57).

### Bait-prey experiments using immobilized AnfO and AnfO domains

Strep-tagged bait proteins were immobilized on gravity flow columns prepared with 500 µL Strep-Tactin®XT 4Flow® resin (IBA Lifesciences Gottingen, Germany). Resin was saturated with the bait protein, and excess bait protein removed by washing with anoxic buffer A (50 mM Tris, 2 mM dithionite, pH 8) exhaustively sparged with Ar and degassed. All manipulations were performed inside a glove box (Coy Laboratory Products, Grass Lake, MI) having an anoxic atmosphere composed by 95% nitrogen and 5% hydrogen. Strep-Tactin resin having no bound bait protein was included as a negative control in all experiments. Cell paste of *A. vinelandii* DJ2239 grown in the presence of 5 µM sodium metavanadate (NaVO_3_) (Sigma-Aldrich, St. Louis, USA) in the culture medium, *A. vinelandii* DJ2520, or *E. coli* BL21 cells expressing the plasmid pDB2343, was resuspended in buffer A and extracts prepared as previously described (18, 52). Before application to the gravity affinity columns, cell extracts were passed over several 5 mL Strep-Tactin columns to remove endogenous biotin-binding proteins such as acetate carboxylase and pyruvate carboxylase. Extract from approximately 25 g of cells for individual experiments was loaded onto a 500 µL gravity column containing an immobilized bait protein and washed with 4-10 column volumes of buffer B (50 mM Tris, 350mM NaCl, 2 mM dithionite, pH 8) until the effluent was clear. Samples were eluted using buffer B containing 50 mM biotin (Sigma-Aldrich, Saint Louis, MO). Eluted samples were analyzed by SDS-PAGE. Captured proteins displayed by SDS-PAGE were identified by MALDI-TOF/TOF analysis performed by the Virginia Tech Mass Spectrometry facility.

### Binding of FeMo-cofactor to AnfO, NifX and NafY

FeMo-cofactor was isolated from MoFe protein purified from DJ1141 cell paste following the reported protocol (35). Thiophenol (PhSH) and aqueous dithionite were added to the as-isolated FeMo-cofactor for final concentrations of 1 mM and 2 mM, respectively. FeMo-cofactor was then added dropwise while stirring to an aqueous solution containing 1.1 equivalents of AnfO, NifX, or NafY, 10% glycerol, 25 mM HEPES (pH 7.5), and 2 mM dithionite. Addition of FeMo-cofactor to the solution results in 100-fold dilution of the N-methylformamide (final concentration: 1% v/v). After 5 minutes of incubation without stirring, the solution was concentrated in an Amicon stirred cell (10 kDa filter for FL AnfO, NifX, and NafY, and 3 kDa filter for the C-domain AnfO).

### Preparing EPR Samples of Phenotypic Test Strains

Purified FeFe protein samples were concentrated to 0.1 mL with a 10 kDa Amicon Ultra-0.5 Centrifugal Filter at 14,000 × g. The solutions were buffer exchanged using Micro Bio-Spin P-6 gel columns that had been equilibrated with 10% glycerol, 25 mM HEPES, pH 7.5 to remove biotin. 2 mM dithionite (final concentration) was then added to generate the EPR samples. EPR spectra were recorded on a Bruker EMX spectrometer at 9.37 GHz as frozen solutions. EPR samples were prepared in an anaerobic glove box with an N_2_ atmosphere and an O_2_ level of <2.5 ppm.

### Data availability

The data supporting the results of this work are available within the article and its supplemental material.

## Supporting information

This article contains supporting information.

## Acknowledgments and funding.

A.P.-G. is a recipient of a Ramón y Cajal Grant (RYC2021-031246-I) from Ministerio de Ciencia e Innovacion (Spain). A.P.-G. was also supported by a Mobility Grant from Programa Propio of Universidad Politécnica de Madrid. Additional support was provided by the National Institutes of Health (GM141203 to D.L.M.S), the Camille and Henry Dreyfus Foundation (TC-22-005 to D.L.M.S.), the Alfred P. Sloan Foundation (FG-2022-18423 to D.L.M.S.), and the MIT Undergraduate Research Opportunities Program (to J.L., with support from the Class of 1973 and Ralph L. Evans (1948) funds) and the U.S. Department of Energy (DE-SC0010834 DRD). Support for the ICP-MS instrument at MIT was provided by a core center grant (P30-ES002109) from the National Institute of Environmental Health Sciences, NIH. This work was supported, in whole or in part, by the Bill & Melinda Gates Foundation BNF Cereals Phase III (INV-005889). Under the grant conditions of the Foundation, a Creative Commons Attribution 4.0 Generic License has already been assigned to the Author Accepted Manuscript version that might arise from this submission.

## Conflict of interest

The authors declare that they have no conflicts of interest with the contents of this article.

## Notes

### Competing Interest Statement

The authors have declared no competing interest.

