## Supporting figures for "Molecular sorting of nitrogenase catalytic cofactors"

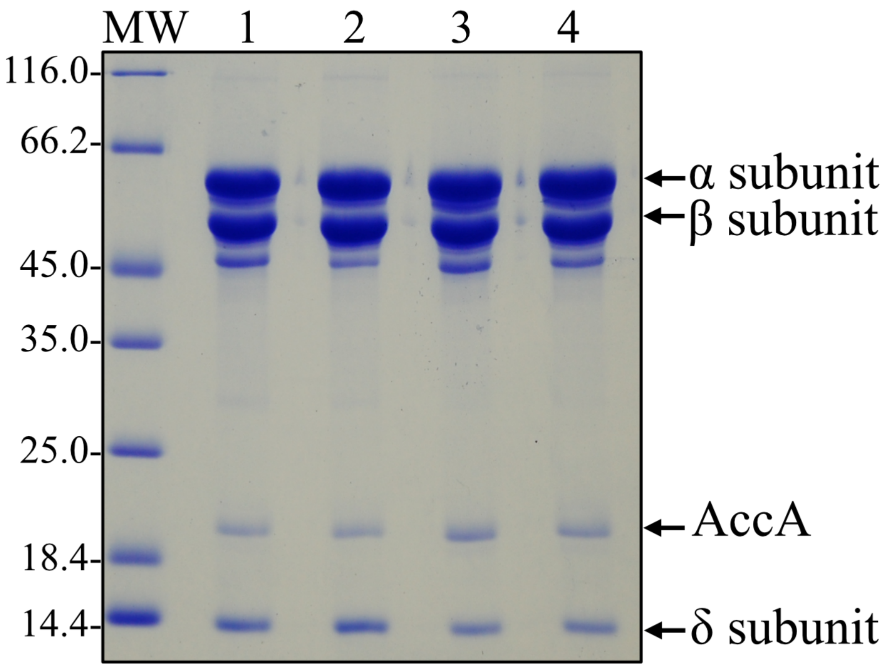


**Figure S1. Coomassie blue stained SDS-PAGE of FeFe proteins purified from different strains and growth conditions**. Lane 1: FeFe protein isolated from DJ2560 (*anfO*$^+$ *nifEN*$^+$); lane 2: FeFe protein isolated from DJ2821 (Δ*anfO nifEN*$^+$); lane 3: FeFe protein isolated from DJ2821 (Δ*anfO nifEN*$^+$) grown in the presence of 1 μM Mo; lane 4: FeFe protein isolated from DJ2831 (Δ*anfO* Δ*nifEN*) grown in the presence of 1 μM Mo. Refer to Tables S1 and S2 for a complete genotypic description of strains. "MW": standard indicating the molecular weight in kDa. Protein identities are indicated by the arrows. Note that the δ subunit is associated with the FeFe protein α- and β-subunits in all the samples analyzed. AccA is the biotin-binding subunit of acetate carboxylase which is unrelated to the FeFe protein but is also isolated from *A. vinelandii* crude extracts using the Strep-Tactin based affinity purification method. The catalytic properties of these isolated FeFe proteins are shown in Table 1.

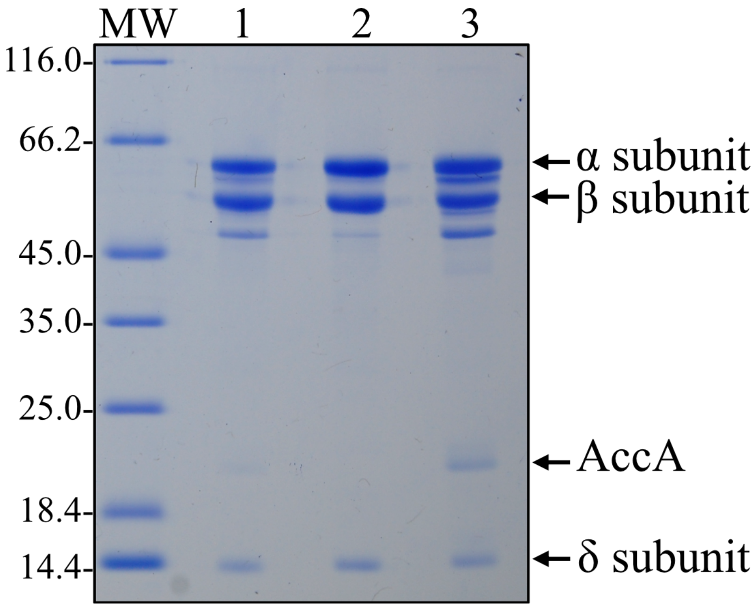


**Figure S2. Coomassie blue stained SDS-PAGE of FeFe proteins containing FeFe-, FeV-, or FeMo- cofactor.** Lane 1: FeFe protein containing FeFe-cofactor from DJ2241 (*anfO*$^{+}$) grown in the absence of Mo and V; lane 2: FeFe protein containing FeV-cofactor from DJ2290 (Δ*anfO vnfEN*$^{+}$) grown in the presence of 5μM V; lane 3: FeFe protein contain FeMo-cofactor from DJ2821 (Δ*anfO nifEN*$^{+}$) grown in the presence of 1 μM Mo. Refer to Tables S1 and S2 for a complete genotypic description of strains. "MW": standard indicating the molecular weight in kDa. Protein identities are indicated by the arrows. Note the δ-subunit is associated with the FeFe protein αβ-subunits in all the samples analyzed.

**A**

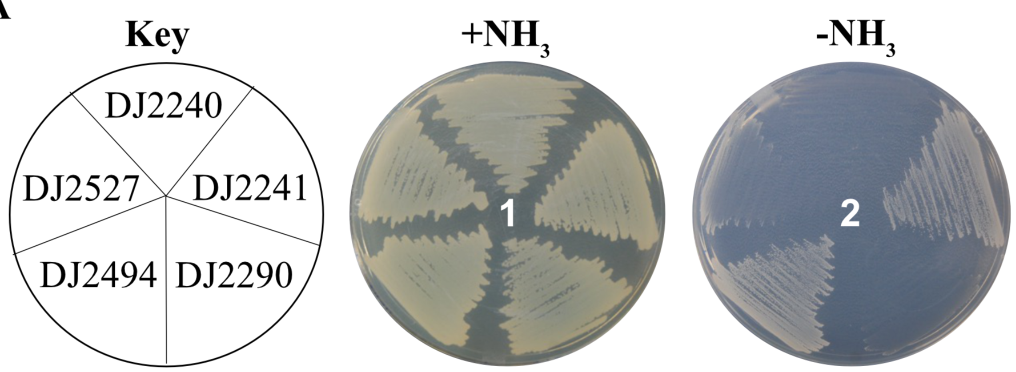


**B**

| Strain | Relevant genotype |
|---|---|
| DJ2240 | Δ*nifDK* Δ*vnfDGK* Δ*anfDGK* |
| DJ2241 | Δ*nifDK* Δ*vnfDGK* *anfD*$^{S\text{-}TAG}$ |
| DJ2290 | Δ*nifDK* Δ*vnfDGK* *anfD*$^{S\text{-}TAG}$ Δ*anfO* |
| DJ2494 | Δ*nifDK* Δ*vnfDGK* *anfO*$^{S\text{-}TAG\,(C\text{-}terminal)}$ |
| DJ2527 | Δ*nifDK* Δ*vnfDGK* *anfO*$^{S\text{-}TAG\,(N\text{-}terminal)}$ |

**Figure S3. Phenotypic characterization of *Azotobacter vinelandii* strains containing a Strep-tag at the N- or C- terminus of AnfO.** A) Strains requiring Fe-only nitrogenase for diazotrophic growth were cultured on Burk's medium agar plates containing a fixed nitrogen source (plate 1) or under diazotrophic conditions (plate 2). Strains were cultured at 30 ºC for 5 days. B) Relevant genes inactivated for each strain. A complete genotypic description of each strain is indicated in Tables S1 and S2. The key observation is that a Strep-tag ("S-TAG") placed at the C-terminus of AnfO (DJ2494) does not impair the *in vivo* function of the protein, while placement of a Strep-tag at the N-terminus of AnfO (DJ2527) or the deletion of a C-terminal portion of AnfO (DJ2290, also DJ2821 shown in Fig. 3) inactivates the *in vivo* function of AnfO based on the diazotrophic growth phenotypes of the corresponding strains.

A

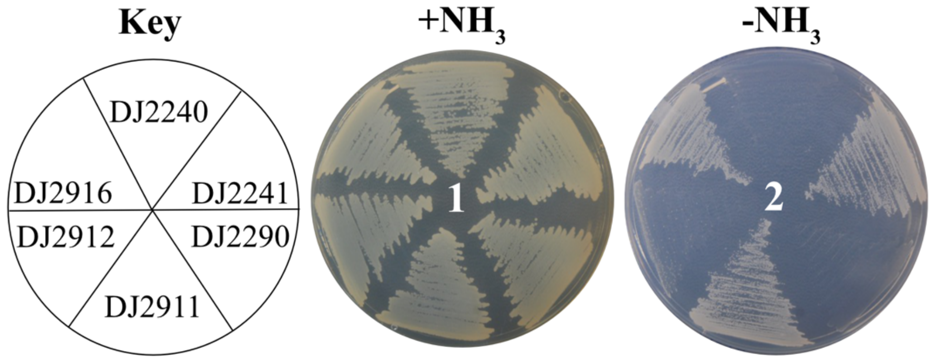


B

| Strain | Relevant genes inactivated |
|---|---|
| DJ2240 | *nifDK, vnfDGK, anfDGK* |
| DJ2241 | *nifDK, vnfDGK* |
| DJ2290 | *nifDK, vnfDGK, anfO* |
| DJ2911 | *nifDK, vnfDGK,* $anfO^{C159A}$ |
| DJ2912 | *nifDK, vnfDGK,* $anfO^{H203L}$ |
| DJ2916 | *nifDK, vnfDGK,* $anfO^{C201A}$ |

**Figure S4. Phenotypic characterization of *Azotobacter vinelandii* strains containing amino acid substitutions within AnfO.** A) Strains requiring Fe-only nitrogenase for diazotrophic growth were cultured on Burk's medium agar plates containing a fixed nitrogen source (plate 1) or under diazotrophic conditions (plate 2). Strains were cultured at 30 ºC for 5 days. B) Relevant genes inactivated for each strain and specific amino acid substitutions of AnfO are indicated. A complete genotypic description of each strain is indicated in Tables S1 and S2. AnfO produced by DJ2911 has the conserved $Cys^{195}$ residue substituted by Ala, DJ2912 has the conserved AnfO $His^{203}$ residue substitute by Leu, and DJ2916 has the conserved AnfO $Cys^{201}$ residue substituted by Ala.

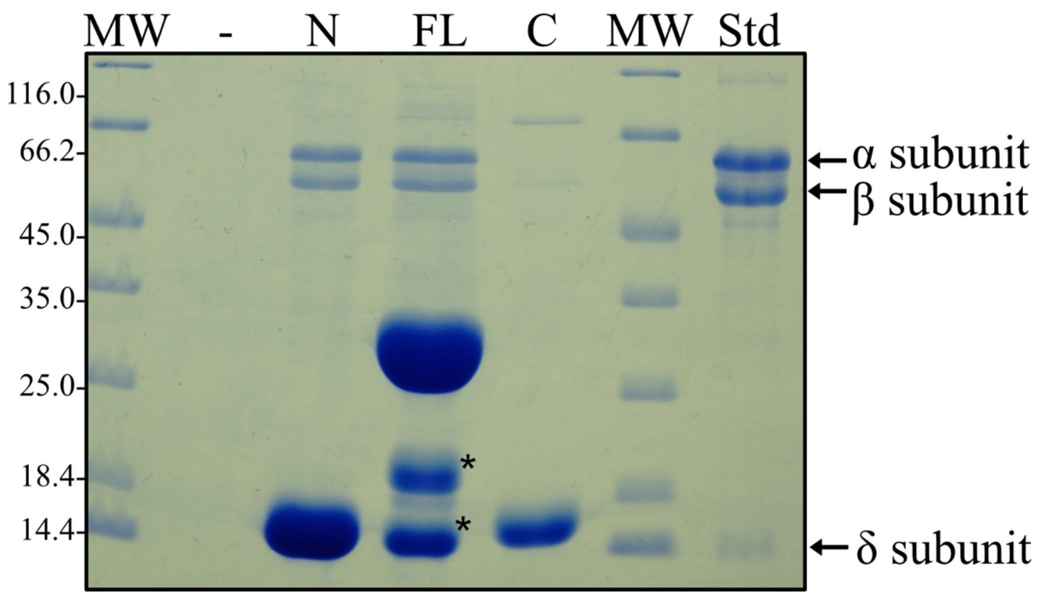


**Figure S5. The apo-FeFe protein interacts with the N-terminal domain of AnfO**. Strep-tagged versions of purified full-length AnfO ("FL"), its N-terminal domain ("N"), or its C-terminal domain ("C") produced in *E. coli* were used to saturate individual Strep-Tactin columns and washed with buffer B as described in Experimental Procedures. Cell extract of DJ2520 (Δ*nifB*, Δ*nifDK*, and Δ*vnfDGK*) was then passed over each column and washed with buffer B containing no biotin, followed by elution with buffer B containing 50 mM biotin. Because DJ2520 carries a *nifB* deletion the FeFe protein, it does not produce FeFe-cofactor and does not contain the δ-subunit. The Coomassie blue stained SDS-PAGE analysis of the eluted samples is shown. Full-length AnfO and its N-domain capture the apo-FeFe protein α- and β-subunits, whereas the AnfO C-domain did not. The second lane "-" represents a negative control for which no bait protein was immobilized on the Strep-Tactin column. MW = molecular weight standards (kDa), FL = immobilized full-length AnfO used as bait, N = immobilized AnfO N-domain used as bait, C = immobilized AnfO C-domain used as bait, Std = purified FeFe protein standard with the αβδ subunits indicated by arrows. The two smaller proteins indicated by asterisks in the lane labelled "FL" are full-length AnfO cleavage products. The identity of all proteins was confirmed by Mass Spectrometry.

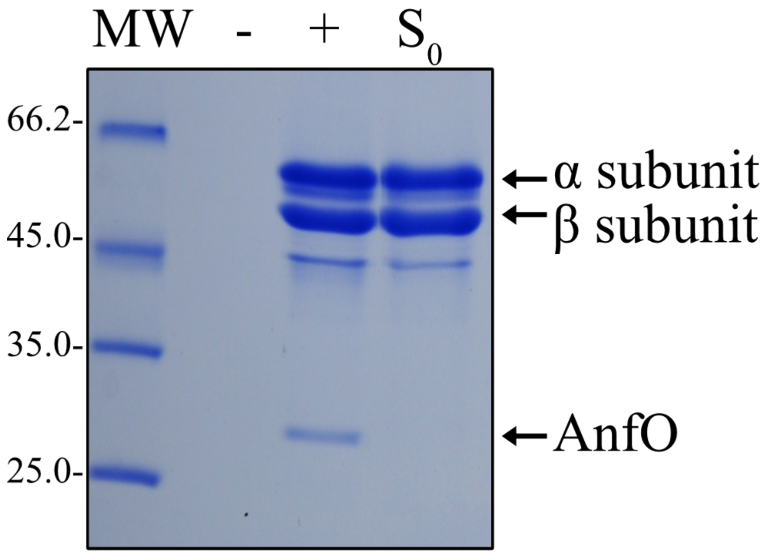


**Figure S6. Immobilized apo-FeFe protein interacts with AnfO.** Purified Strep-tagged apo-FeFe protein produced by *A. vinelandii* strain DJ2245 was used to saturate individual Strep-Tactin columns and washed with buffer B as described in Experimental Procedures. Cell extract of *E. coli* BL21 cells expressing a plasmid containing a non-tagged version of full-length AnfO (pDB2343) was then passed over each column and washed with buffer B containing no biotin, followed by elution with buffer B containing 50 mM biotin. Coomassie blue stained SDS-PAGE analysis of the eluted samples is shown. The apo-FeFe protein α- and β-subunits capture full-length AnfO as shown in lane "+". Lane "$S_0$" shows an immobilized apo-FeFe protein sample for which no *E. coli* cell extract was passed over the column. Lane "-" represents a negative control for which no bait protein was immobilized on the Strep-Tactin column. MW = molecular weight standards (kDa). Protein identities are indicated by arrows. Identity of full-length AnfO was confirmed by Mass Spectrometry.
