## Supporting tables for "Molecular sorting of nitrogenase catalytic cofactors"

**Table S1. List of *Azotobacter vinelandii* strains.**

| Strain | Genotype |
|---|---|
| DJ2239 | *ΔnifDK*, *vnfDGK*::$sm^R$,$rif^R$, Δ42kb[a] |
| DJ2240 | *ΔnifDK*, *vnfDGK*::$sm^R$, *anfDGK*::$km^R$, $rif^R$, Δ42kb[a] |
| DJ2241 | *ΔnifDK*, *vnfDGK*::$sm^R$, $anfD^{S\text{-}TAG}$, $rif^R$, Δ42kb[a] |
| DJ2245 | *ΔnifDK*, *vnfDGK*::$sm^R$, Δ*nifB*::$km^R$, $anfD^{S\text{-}TAG}$, $rif^R$, Δ42kb[a] |
| DJ2290 | *ΔnifDK*, *vnfDGK*::$sm^R$, Δ*anfO*, $anfD^{S\text{-}TAG}$, $rif^R$, $tet^R$, Δ42kb[a] |
| DJ2494 | *ΔnifDK*, *vnfDGK*::$sm^R$, $anfO^{S\text{-}TAG\ (\text{C-term})}$, $rif^R$, $tet^R$, Δ42kb[a] |
| DJ2520 | *ΔnifDK*, *vnfDGK*::$sm^R$, Δ*nifB*, $rif^R$, Δ42kb[a] |
| DJ2527 | *ΔnifDK*, *vnfDGK*::$sm^R$, $anfO^{S\text{-}TAG\ (\text{N-term})}$, $rif^R$, $tet^R$, Δ42kb[a] |
| DJ2560 | *ΔnifDK*, *vnfDGK*::$sm^R$, *vnfE*::$gm^R$, Δ*modE1*[a], $anfD^{S\text{-}TAG}$, $rif^R$ |
| DJ2821 | *ΔnifDK*, *vnfDGK*::$sm^R$, *vnfE*::$gm^R$, Δ*modE1*[a], Δ*anfO*, $anfD^{S\text{-}TAG}$, $rif^R$ |
| DJ2831 | *ΔnifDK*, *vnfDGK*::$sm^R$, *vnfE*::$gm^R$, Δ*modE1*[a], Δ*anfO*, *nifE*::$km^R$, $anfD^{S\text{-}TAG}$, $rif^R$ |
| DJ2911 | *ΔnifDK*, *vnfDGK*::$sm^R$, $anfD^{S\text{-}TAG}$, $anfO^{C159A}$, Δ42kb[a] |
| DJ2912 | *ΔnifDK*, *vnfDGK*::$sm^R$, $anfD^{S\text{-}TAG}$, $anfO^{H203L}$, Δ42kb[a] |
| DJ2916 | *ΔnifDK*, *vnfDGK*::$sm^R$, $anfD^{S\text{-}TAG}$, $anfO^{C201A}$, Δ42kb[a] |

$anfD^{S\text{-}TAG}$: Strep-tag is placed at the C-terminal of *anfD;* $rif^R$: rifampicin; $sm^R$: streptomycin; $km^R$: kanamycin; $tet^R$: tetracycline; $gm^R$: gentamycin.

Location of residues removed and/or placement of insertions are indicated in Table S2.

[a] W-tolerance is the result of a Δ42kbp in a genomic deletion required for Mo acquisition or the Δ*modE1* whose product is involved in regulating Mo acquisition and Mo-dependent repression of *anf* gene expression.

**Table S2. List of plasmids used for the construction of *Azotobacter vinelandii* strains**.
Location of residues removed and/or placement of insertions are indicated. Nomenclature corresponds to the genotype shown in Supplemental Table 1. S-TAG: Strep-tag (ASWSHPQFEK); $km^R$: kanamycin; $sm^R$: streptomycin; $gm^R$: gentamycin.

| **Plasmid** | **Deletion/Insertion** | **Residues Removed/ Insertion Location** |
|---|---|---|
| pDB33 | Δ*nifDK* | $NifD^{103}$ - $NifK^{308}$ |
| pDB161 | Δ*nifB* | $NifB^{60\text{-}307}$ |
| pDB218 | *nifB*::$km^R$ | $NifB^{60\text{-}307}$ |
| pDB259 | *nifE*::$km^R$ | $NifE^{15\text{-}261}$ |
| pDB2134 | *anfDGK*::$km^R$: | $AnfD^{204}$ - $AnfK^{148}$ |
| pDB2139 | *vnfDGK*::$sm^R$ | $VnfD^{271}$ - $VnfK^{202}$ |
| pDB2158 | $anfD^{S\text{-}TAG}$ | $AnfD^{518}$ |
| pDB2200 | *vnfE*::$gm^R$ | $VnfE^{91}$ |
| pDB2224 | Δ*anfO* | $AnfO^{179\text{-}224}$ |
| pDB2265 | Δ*modE1* | $ModE1^{151\text{-}215}$ |
| pDB2355 | $anfO^{S\text{-}TAG}$ (C-term) | $AnfO^{245}$ |
| pDB2395 | $anfO^{S\text{-}TAG}$ (N-term) | $AnfO^{1}$ |
| pDB2611 | | $AnfO^{C159A}$ |
| pDB2612 | | $AnfO^{C201A}$ |
| pDB2613 | | $AnfO^{H203L}$ |

**Table S3. Plasmids used for heterologous expression of AnfO and the N- and C-domains in *Escherichia coli* BL21(DE3) competent cells.**

| Plasmid | Description |
|---|---|
| pDB2343 | For purification of the non-tagged, full-length AnfO (residues 1-245). |
| pDB2418 | For purification of the full-length AnfO (residues 1-245). A Strep-tag (ASWSHPQFEK) is located after residue 245. |
| pDB2526 | For purification of the C-terminal domain of AnfO (residues 138-245). A TwinStrep-tag (ASWSHPQFEKGGGSGGGSGGSAWSHPQFEKAS) is located before residue 138. |
| pDB2554 | For purification of the N-terminal domain of AnfO (residues 1-132). A Strep-tag (ASWSHPQFEK) is located after residue 132. |

**Table S4. Quantification of FeMo-cofactor binding to AnfO based ICP-MS metal and BCA protein assays.**

| Sample | Fe | Mo | AnfO | [Mo] : [AnfO] |
|---|---|---|---|---|
| 1 | 8.4 | 1 | 1.6 | 0.62 |
| 2 | 7.1 | 1 | 2.3 | 0.44 |
| 3 | 6.6 | 1 | 1.9 | 0.53 |
| 4 | 11 | 1 | 2.2 | 0.45 |
| **Combined** | **7.1** | **1** | **2.0** | **0.51** |

The “Combined” sample was prepared by mixing samples 1-4 and concentration using a stirred cell concentrator. These data show that repeated generation of AnfO containing FeMo-cofactor results in an [Fe] : [Mo] ratio close to 7:1, as is expected for a FeMo-cofactor containing protein. Some occurrences of higher [Fe] : [Mo] ratio have been observed (notably sample 4), which we attribute to cluster degradation. Additionally, the [Mo] : [AnfO] ratio, which we use as a proxy for the [FeMo-cofactor] : [AnfO] ratio, consistently resides in the range of $0.5 \pm 0.1$. At this time, we cannot distinguish between weak FeMo-cofactor binding to AnfO (and a native 1:1 cofactor:AnfO ratio) or a native 1:2 cofactor:AnfO ratio.
